# Local biodiversity change reflects interactions among changing abundance, evenness and richness

**DOI:** 10.1101/2021.08.29.458087

**Authors:** Shane A. Blowes, Gergana N. Daskalova, Maria Dornelas, Thore Engel, Nicholas J. Gotelli, Anne E. Magurran, Inês S. Martins, Brian McGill, Daniel J. McGlinn, Alban Sagouis, Hideyasu Shimadzu, Sarah R. Supp, Jonathan M. Chase

**Affiliations:** German Centre for Integrative Biodiversity Research (iDiv) Halle-Jena-Leipzig, Germany; Department of Computer Science, Martin Luther University Halle-Wittenberg, Halle (Salle), Germany.; School of GeoSciences, University of Edinburgh, Scotland EH9 3FF.; Centre for Biological Diversity, University of St Andrews, KY16 9TH.; German Centre for Integrative Biodiversity Research (iDiv) Halle-Jena-Leipzig, Germany; Department of Computer Science, Martin Luther University Halle-Wittenberg, Halle (Salle), Germany.; Department of Biology, University of Vermont, Burlington, VT 05405 USA.; Leverhulme Centre for Anthropocene Biodiversity and Department of Biology, University of York, York YO10 5DD, UK; Centre for Biological Diversity, University of St Andrews, KY16 9TH.; School of Biology and Ecology and Mitchell Center for Sustainability Solutions, University of Maine; Orono, ME.; Department of Biology, College of Charleston, Charleston, SC.; Department of Mathematical Sciences, Loughborough University, LE11 3TU, UK; Graduate School of Public Health, Teikyo University, Tokyo, Japan.; Data Analytics Program, Denison University, Granville, OH 43023 USA.

**Keywords:** biodiversity change, abundance, evenness, species richness, rarefaction

## Abstract

Biodiversity metrics often integrate data on the presence and abundance of multiple species. Yet understanding covariation of changes to the numbers of individuals, the evenness of species’ relative abundances, and the total number of species remains limited. Using individual-based rarefaction curves, we introduce a conceptual framework to understand how expected positive relationships among changes in abundance, evenness and richness arise, and how they can break down. We then examined interdependencies between changes in abundance, evenness and richness in more than 1100 assemblages sampled either through time or across space. As predicted, richness changes were greatest when abundance and evenness changed in the same direction, and countervailing changes in abundance and evenness acted to constrain the magnitude of changes in species richness. Site-to-site changes in abundance, evenness, and richness were often decoupled, and pairwise relationships between changes in these components across assemblages were weak. In contrast, changes in species richness and relative abundance were strongly correlated for assemblages varying through time. Temporal changes in local biodiversity showed greater inertia and stronger relationships between the component changes when compared to site-to-site variation. Local variation in assemblage diversity was rarely due to a passive sample from a more or less static species abundance distribution. Instead, changing species relative abundances often dominated local variation in diversity. Moreover, how changing relative abundances combined with changes to total abundance frequently determined the magnitude of richness changes. Embracing the interdependencies between changing abundance, evenness and richness can provide new information for better understanding biodiversity change in the Anthropocene.

## Introduction

Measures of biodiversity and its change are frequently used to determine the magnitude, direction and pace of ecosystem modifications (Diaz et al. 2019). Descriptions of the distribution and abundance of organisms are central to basic ecological research (Krebs 1972, Andrewartha & Birch 1984), and biodiversity (as applied to taxonomic or species diversity) is a multifaceted concept that combines information on the distribution and abundance of multiple species. There have been numerous such aggregate metrics proposed to quantify different aspects of biodiversity (e.g., Hill 1973, Gotelli and Colwell 2001, Magurran & McGill 2011, Chao and Jost 2012), most of which depend on sample effort and scale (Rosenzweig 1995, Whittaker et al. 2001, McGill 2011a, Chase & Knight 2013). This inherent complexity poses significant challenges for quantitative syntheses of temporal and spatial variation in biodiversity.

Here, we argue that in order to understand biodiversity change, it is critical to look beyond aggregate metrics in isolation. Specifically, we need to examine how changes in the key components that lead to aggregate metrics interact and combine with each other. Currently, different components that underlie biodiversity represent important, but are largely viewed as independent lines of evidence for contemporary biodiversity change. For example, one body of work is focused on quantifying total abundances of all species within assemblages, such as recent work documenting declines of birds (e.g., Rosenberg et al. 2019) and insects (Wagner 2020). A related body of work focuses instead on population-level ‘winners’ and ‘losers’ within assemblages (McKinney & Lockwood 2000), species with increasing and decreasing trends in occupancy or abundance (e.g., WWF 2020). However, changes in the means of these metrics do not capture the variability and nuance in either the assemblage- and population-level abundance trends across space and among taxa (e.g., van Klink et al. 2020, Daskalova et al. 2020a, Leung et al. 2020). Moreover, trends in total assemblage abundance provide only one window into potentially complex changes that could be occurring in other components of biodiversity, and while changes to species relative abundances do influence biodiversity metrics, population-level trends themselves cannot be used to calculate assemblage-level (i.e., biodiversity) metrics (e.g., Dornelas et al. 2019). Even when quantifying the same response metric (i.e., species richness), syntheses of biodiversity change in space due to differential land use conditions have found declines of species richness (e.g., Newbold et al. 2015, 2018), whereas analyses of time series have shown that, on average, species richness increases roughly offset decreases (Dornelas et al. 2014, Blowes et al. 2019), despite often significant modifications to climate and habitat (Antão et al. 2020a, Daskalova et al. 2020b). Finally, the prediction that human activities likely impact species relative abundances more frequently than species occurrences (Chapin et al. 2000), has not resulted in a strong focus on assemblage-level evenness in existing syntheses (but see, e.g., Crowder et al. 2010, Zhang et al. 2012, Jones & Magurran 2018). Importantly, changes in all these components -- abundance, evenness, and richness -- contribute to variation in biodiversity. Yet little is known about how components are changing in combination within assemblages, and whether certain combinations act to constrain observed biodiversity change.

Species origination (speciation plus colonization) and extinction are the most fundamental processes for biodiversity dynamics (Storch et al. 2021). These processes combine with productivity (Currie 1991, Mittelbach et al. 2001), disturbance frequency and intensity (Connell 1978, Miller et al. 2011), historical (Latham & Ricklefs 1993) and biogeographic factors (e.g., Kreft et al. 2008), land use modifications (Newbold et al. 2015), and climate change (Antão et al. 2020a) to drive variation in biodiversity. In turn, these processes and drivers combine such that local scale measures of biodiversity estimated from a given (local) sample depends largely on two components (see e.g., He and Legendre 2002, McGill 2011a). First, the total number of individuals (Fisher et al. 1943, Preston 1962), whereby fewer individuals are expected to (non-linearly) lead to fewer species. Second, the total number of species and their relative abundances within the regional species pool (i.e., the set of all potential colonizing species in a region), which we refer to as the Species Abundance Distribution (SAD; McGill et al. 2007). Local samples will have lower species richness when the species pool has relatively few highly abundant species and many rare species (i.e., less even SADs), compared to samples from a species pool of the equivalent size with more equitable species abundances. Moreover, variation or changes in local environmental conditions can alter local patterns of species relative abundances. Whenever two or more samples across space or time differ in the total number of individuals, the shape of the SAD, or both, there will be changes in most metrics of biodiversity. However, changes in abundance and the SAD are not always correlated, and, when decoupled, the magnitude and direction of change in derived biodiversity metrics can differ considerably.

Variation in the total number of individuals is a long-standing, first-order explanation of variation in richness (Fisher et al. 1943, Coleman et al. 1982, Srivastava & Lawton 1998, Gaston 2000, Scheiner & Willig 2005, Storch et al. 2018). In the context of species-area relationships, this has been termed the ‘passive sampling hypothesis’ (Coleman et al. 1982), and as local assemblages increase in size they are expected to include more species from the regional pool due to sampling processes alone. Larger (Connor & McCoy 1979) or more productive (Wright 1983) areas are also predicted to have more species driven again by increased numbers of individuals, but in these cases associated with processes other than sampling, such as decreased likelihood of species extinction with increased population sizes (Preston 1962, Wright 1983, Srivastava & Lawton 1998), and commonly termed the ‘more individuals hypothesis’ (Srivastava & Lawton 1998). Anthropogenic drivers can also influence the number of individuals in assemblages (e.g., via eutrophication, exploitation, harvesting or land clearing), potentially impacting biodiversity due to changes to the total number of individuals (Newbold et al. 2015, Blowes et al. 2020). If biodiversity varies primarily via changes in the numbers of individuals, positive relationships between altered numbers of individuals and species richness changes are expected. In such cases, however, other metrics of species diversity that control for variation in numbers of individuals, such as species richness expected for a given number of individuals— known as rarefied richness—should be relatively unchanged.

Changes to the shape of the SAD can drive variation in biodiversity through time or space. For example, co-occurrence and coexistence of species can be altered by changes to resource diversity (MacArthur 1965), environmental or habitat heterogeneity (Tilman 1982, Shmida & Wilson 1985), interspecific interactions (e.g., keystone predation; Paine 1974, Menge et al. 2020), biological invasions (Vilà et al. 2011), and external perturbations (Hughes et al. 2007). Alterations to any of these features can change biodiversity by changing species’ relative abundances and the size of the species pool (via species additions or subtractions).

Anthropogenic factors can also favor some species and disfavor others, potentially altering the relative abundance of species (e.g., due to selective exploitation; Blowes et al. 2020), or the size of the species pool (e.g., species with large ranges replacing those with small ranges, Newbold et al. 2018). In such cases, biodiversity change will be characterized by positive relationships between species richness change and changes in metrics sensitive to relative abundance, such as rarefied richness, evenness and diversity metrics sensitive to species relative abundances.

Changing components of biodiversity can covary in different and informative ways. Yet, to date, there has been little exploration of this covariation in time or space, nor of the theoretical linkages. For example, whether total abundances and the evenness of species’ relative abundances change in similar or decoupled ways, and how this influences biodiversity change is largely unknown. However, syntheses of relationships between different biodiversity metrics, which can reflect different combinations of component changes, have typically found relationships to be weak. For example, Stirling and Wilsey (2001) showed that although strong positive correlations between species richness, diversity and evenness metrics were expected from a neutral model (Caswell 1976), there was considerable variation in the strength, and even the sign of the relationships in 323 empirical comparisons. Similarly, Soininen et al. (2012) examined temporal (*n* = 212) and spatial variation (*n* = 17) in aquatic datasets, and again found considerable heterogeneity in the relationship between richness and evenness. Using data from 91 assemblages, McGill (2011b) concluded that most biodiversity metrics align with three axes of empirical variation (total abundance, evenness and richness); components subsequently shown to be relatively uncorrelated across space for a subset of 37 of the 91 assemblages (Chase et al. 2018). Collectively, these studies suggest that static biodiversity estimates are multidimensional, and that different metrics can covary or be unrelated.

Where ecologists have quantified variation in multiple components of local diversity, the focus has typically been on averages across assemblages, with each component treated as a separate, independent response. For example, analyses of the local assemblages documented by the BioTIME database (Dornelas et al. 2018) show that numbers of individuals, species richness, and dominance (quantified as the relative abundance of the most numerically dominant species, and conceptually the complement of evenness) are highly variable among datasets, but on average, have no directional trend (Dornelas et al. 2014, Jones & Magurran 2018, Blowes et al. 2019). On the other hand, analyses of the PREDICTS database (Hudson et al. 2017) documenting spatial contrasts between assemblages in more pristine habitats with those in different land use categories, show that human-altered habitats frequently have fewer species and often fewer individuals (Newbold et al. 2015, 2020). However, these results describe average changes across assemblages estimated independently, whereas, as we describe in more detail below, component changes are unlikely to be completely independent.

Here, we first provide a conceptual overview of how changes in the main components underlying local biodiversity (total abundance, evenness and species richness) can combine using individual-based rarefaction curves. Using simplified scenarios with contrasting component changes, we show that the signs (or direction) of changes in total abundance and evenness can combine to determine the magnitude of expected richness changes, and whether positive pairwise relationships prevail. We then empirically assess interdependencies between abundance, evenness and richness changes using compilations of ecological assemblage data. In the face of natural and anthropogenically-driven environmental variation in time and space, we ask whether changes in the components of local biodiversity show positive relationships (i.e., change in the same direction). Or, alternatively, are component changes sufficiently heterogeneous that variation in biodiversity depends on which of the underlying components (numbers of individuals or the SAD) are changing, and how the different component changes combine?

### Conceptual relationships between changes in total abundance, evenness and richness

Individual-Based Rarefaction (IBR) curves (Hurlbert 1971, Gotelli & Colwell 2001) are well suited for visualizing conceptual relationships among changes in total abundance, evenness and species richness (Fig. 1a, Cayuela et al. 2015, Chase et al. 2018, McGlinn et al. 2019). The end point of the IBR depicts the total number of individuals of all species combined, and variation between assemblages in where the IBR terminates quantifies changes to the number of individuals (ΔN, Fig.1a) and species richness (ΔS, Fig. 1a). The shape (or curvature) of the IBR curve reflects species’ relative abundances and the size of the species pool (i.e., the SAD). We use two parts of the curve to characterize changes in the SAD between assemblages. First, because it is standardized to an equal number of individuals (*n*), changes in rarefied richness, Δ*S*_n_ (Fig.1a), reflects changes to species’ relative abundances only. Second, we use the numbers equivalent (or effective number of species) transformation of the Probability of Interspecific Encounter (PIE, Hurlbert 1971). The PIE is equal to the slope at the base of the rarefaction curve (Olszewski 2004) and represents a metric of evenness that is relatively insensitive to sample effort (more even communities have a higher PIE, Fig. 1a). Transformation of the PIE to the numbers equivalent (*S*_PIE_) aids comparisons to species richness (i.e., Δ*S* and Δ*S*_PIE_ have the same units; Jost 2006). *S*_PIE_ is equal to the inverse of Simpson concentration (Jost 2006), and diversity of order *q* = 2 (Hill 1973, Jost 2007), 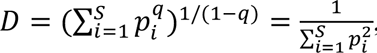 , where *S* is the number of species and *p*_i_ is the proportion of the assemblage represented by species *i*. As a consequence, changes in *S*_PIE_ (Δ*S*_PIE_) are most strongly influenced by the number of abundant or common species in assemblages.

**Figure 1:**
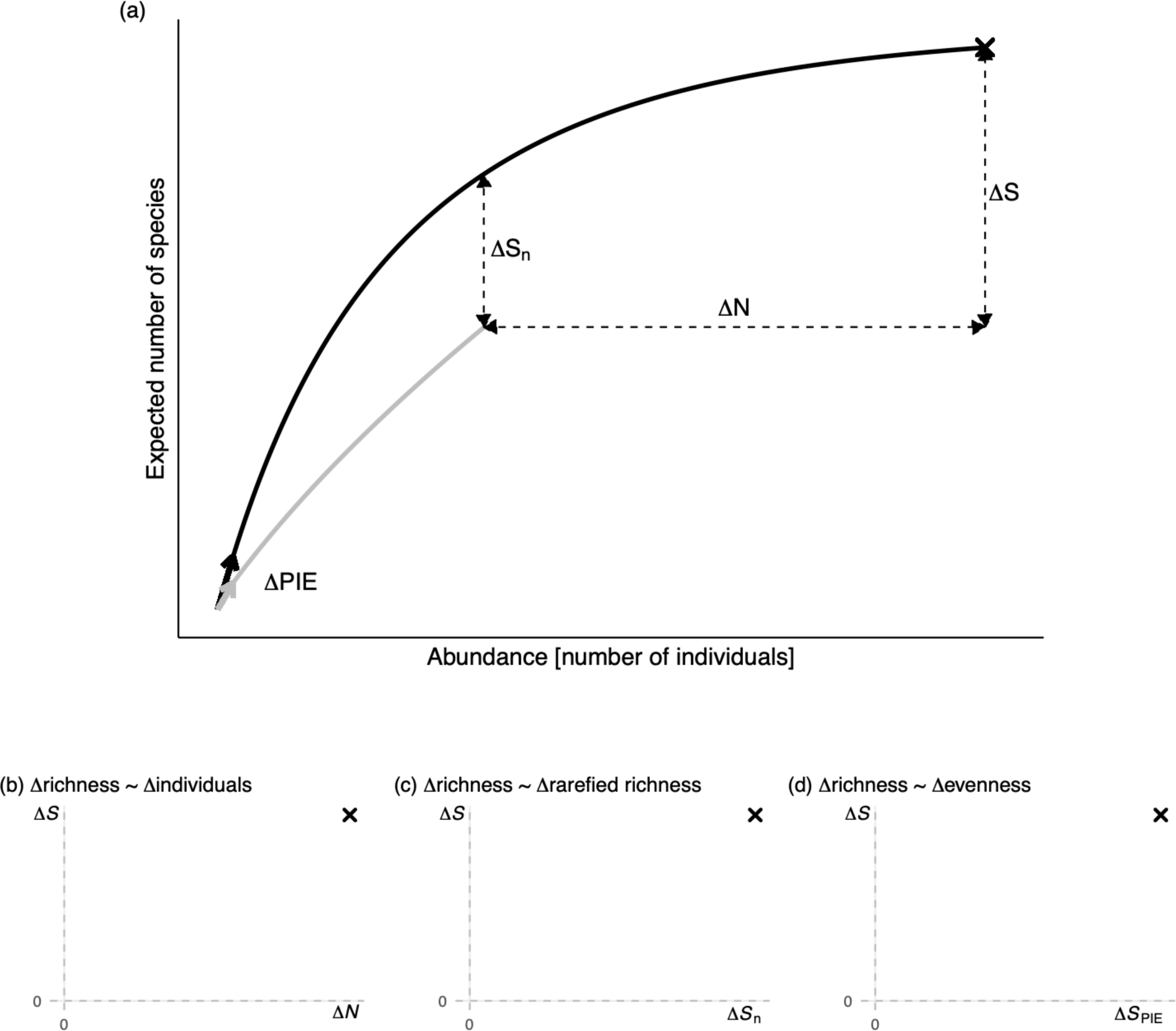
(a) Individual-based rarefaction (IBR) curves for two hypothetical assemblages, showing the four components we use to quantify change (*N*, *S*, *S*_n_, PIE). We visualize relationships between these four components of the IBR curve by plotting: (b) changes in species richness (Δ*S*) as a function of altered numbers of individuals (Δ*N*), (c) changes in species richness (Δ*S*) as a function of changes in rarefied richness (Δ*S*_n_), and (d) changes in species richness (Δ*S*) as a function of the numbers equivalent conversion of the Probability of Interspecific Encounter (*ΔS*_PIE_). We show *Δ*PIE on the figure to illustrate changes of the PIE (it is equivalent to the slope at the base of the curve) with the IBR, but use the numbers equivalent transformation (*ΔS*_PIE_) in all analyses. Points on panels (b, c, and d) show changes between the two hypothetical assemblages, with the reference assemblage depicted by the grey line.

To visualize relationships among changes in total abundance, evenness and richness, we plot pairwise relationships between changes in four components of the IBR. Specifically, changes in species richness (Δ*S*) and total abundance (Δ*N*; Fig. 1b), and species richness and the two metrics sensitive to changes in relative abundance – rarefied richness, Δ*S*_n_ (Fig. 1c), and Δ*S*_PIE_, which we refer to as evenness (Fig. 1d). Changing components with positive relationships (i.e., the same sign) fall into the lower left and upper right quadrants, whereas assemblage changes that fall into the upper left or lower right quadrants of Figs. 1b, c, d reflect a negative relationship between the respective components.

Altered numbers of individuals, but no change to the SAD, can underpin differences in diversity between assemblages. Changes only to the number of individuals being passively sampled from the same underlying SAD (Fig. 2a) result in Δ*S* and Δ*N* being positively related with the same sign (Fig. 2g), whereas Δ*S*_n_ (Fig. 2h) and Δ*S*_PIE_ (Figs. 2i) will be approximately zero (and have a weak or no relationship with Δ*S*). This has been variously referred to in the literature as a sampling effect, the rarefaction effect, and the passive sampling hypothesis (Coleman et al. 1982, Gotelli & Cowell 2001, Palmer et al. 2000).

**Figure 2:**
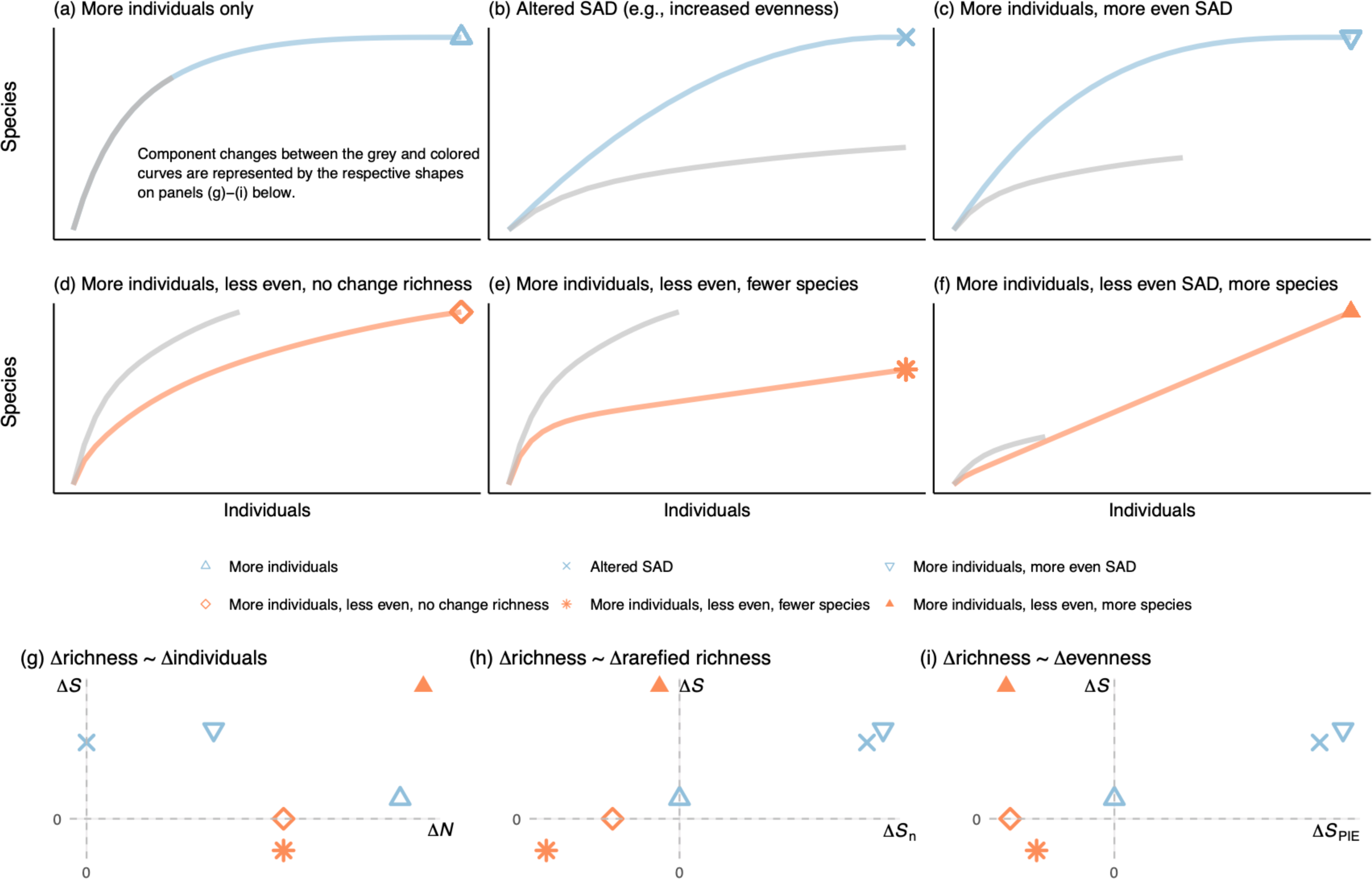
Conceptual illustrations of potential pathways of local assemblage diversity change and corresponding relationships between component changes. Starting from a reference assemblage (depicted with grey rarefaction curves), diversity change can be due to (a) more individuals only, (b) changes to the species abundance distribution only (e.g., increased species pool size or increased evenness), or (c) changes in total abundance and the SAD that result in positive pairwise relationships between Δ*N,* Δ*S*_n_, Δ*S*_PIE_, and Δ*S*. However, if the signs of Δ*N* and Δ*S*_PIE_ differ, their relationships with Δ*S* weaken and species richness can (d) remain static, (e) decrease or (f) increase. We visualize pairwise relationships between component changes for each scenario (i.e., the different shaped symbols) using: (g) changes in species richness as a function of changes to the number of individuals, (h) changes in species richness as a function of changes in rarefied richness, and (i) changes in species richness as a function of changes in evenness.

Changes in species richness (Δ*S*) can also be solely associated with changes to relative abundance (i.e., Δ*N* ≈ 0), which weakens or removes the expectation for a positive relationship between changes in richness and total abundance. For example, changes in species richness can be associated with SAD changes due, for e.g., to increased numbers of common species, increased evenness (He & Legendre 2002), or increases to the size of the species pool (Fig. 2b), which results in a positive relationship between Δ*S* and Δ*S*_PIE_ (Figure 2i). Finally, if total abundance and relative abundance change in the same direction (e.g., more individuals and increased evenness, Fig. 2c), then positive pairwise relationships are expected between changes in abundance, evenness and richness (Figs. 2g, h, i).

In contrast, even if numbers of individuals increase (Δ*N* > 0), expected gains in species richness can be constrained by decreased evenness. For example, opposing changes in total abundance and evenness can potentially result in no change to species richness (Fig. 2d), and no relationship between Δ*S* and Δ*N* (Fig. 2g). Or, if changes to the SAD are sufficiently strong, they can offset any expected gains due to more individuals (Fig. 2e), and result in a negative relationship between Δ*S* and Δ*N* (Fig. 2g). Alternatively, opposing changes to total numbers of individuals and evenness could result in a positive relationship between Δ*S* and Δ*N* if, for example, the effects of more individuals on species richness outweighs that of decreased evenness (Figure 2f).

These simplified scenarios illustrate the potential for interdependencies between component changes. In particular, they show that the signs of changes in total abundance and evenness (i.e., Δ*N* and Δ*S*_PIE_) can strongly influence the magnitude of richness changes, and whether expected positive relationships between changes in abundance, evenness and richness are found. Δ*N* is associated with the IBR curve stretching or contracting along the x-axis, and Δ*S*_PIE_ characterizes changes that flex the curve up or down from the base along the y-axis (Olszewski 2004). When Δ*N* and Δ*S*_PIE_ have the same sign, assemblages are expected to fall into the lower left and upper right quadrants of Figs. 2g-i (i.e., component changes with the same sign and positive pairwise relationships). In contrast, when Δ*N* and Δ*S*_PIE_ have different signs, they can have countervailing effects that constrain richness changes, the strength of their pairwise relationships with Δ*S* will be diminished and potentially reversed, and the likelihood of assemblages falling into the upper left and lower right quadrants of Figs. 2g-i increases (i.e., opposing signs and negative pairwise relationships).

### Empirical relationships among total abundance, evenness and richness

Next, we evaluate empirical relationships by fitting models that allow for correlations between component changes to data from 1125 assemblages. Our goal for the empirical analyses was to examine relationships between changing components in temporal and spatial contexts across a broad range of environmental conditions. We compiled data documenting either temporal or spatial variation of assemblage composition in one of either naturally-varying or perturbed environmental conditions. Temporal variation quantified rates of change (i.e., per year) for each component for an assemblage at a single location through time. Analyses of spatial variation quantified component changes between sites in different land use categories in perturbed environments, or between random pairs of sites in the naturally-varying environment.

Based on our conceptual overview, we expect pairwise relationships between abundance, evenness and richness changes to be generally positive. Changes in species richness are also expected to be largest for assemblages where all pairwise relationships are positive. In contrast, opposing changes in total abundance and evenness (i.e., Δ*N* and Δ*S*_PIE_ have different signs) are expected to constrain changes in species richness. Additionally, if local diversity changes are dominated by altered total abundances and species richness, strong positive relationships between Δ*S* and Δ*N*, but weaker relationships between Δ*S* and Δ*S*_n_, and Δ*S* and Δ*S*_PIE_ should emerge across assemblages. Alternatively, strong relationships between either Δ*S* and Δ*S*_n_ and/or Δ*S* and Δ*S*_PIE_, accompanied by a weaker relationship between Δ*S* and Δ*N,* would indicate that changes to the SAD are the dominant component of local assemblage change.

### Temporal comparisons: natural environmental variation

Temporal changes in natural assemblages at the local scale were quantified using the BioTIME database (Dornelas et al. 2018). Annual rates of change (change per year) for each metric were estimated with models fit to data that documents over 45 thousand species in time series with an average duration of 13 years. Taxonomic groups in our analysis came from surveys in marine, freshwater and terrestrial ecosystems, and included plants (and other producers), invertebrates, fish, amphibians, reptiles, birds, and mammals, as well as several surveys that collected data from multiple taxa. Here, we only used time series that had numerical abundance data available (i.e., studies that recorded counts of the number of individuals for each species in an assemblage), and our analysis included 288 studies. Locations sampled in the BioTIME database document places with varying degrees of anthropogenic environmental change, but do not include manipulated assemblages or before-after-control-impact studies (Dornelas et al. 2018).

Accordingly, we contrast the environmental variation sampled by BioTIME with assemblage time series that experienced documented perturbations (see *Temporal comparisons: experimental or natural perturbations*).

To quantify changes at the local scale within BioTIME, studies with large extents were broken up into smaller cell-level time series, while still maintaining the integrity of individual studies (i.e., different studies were not combined, even when samples were collected in the same grid cell). We used sample-based rarefaction (Gotelli & Colwell 2001) to standardize the number of samples per year for each time series (see Blowes et al. 2019 for details). For the calculation of rarefied richness (*S*_n_), the minimum total number of individuals was determined for each time series, and set as the target *n* for which expected richness was calculated; cell-level time series where *n* < 5 were discarded. This process resulted in 42,604 cell-level time series from the 288 studies, and we focus on the study-level estimates of change in our results and discussion.

### Temporal comparisons: experimental or natural perturbations

To complement the environmental variation sampled by the BioTIME database, we searched for time series data with either experimental or natural perturbations. Specifically, we queried the U.S. LTER network using the Data Portal of the Environmental Data Initiative (https://portal.edirepository.org/nis/home.jsp) with the search terms ‘experiment’ and ‘time’ and ‘abundance’. Records returned were checked to confirm that samples documented assemblages of similar species collected with the same methodology, and following data standardization (i.e., minimum of five individuals per sample, and standardization of sample effort through time), our analysis included 11 studies (see Appendix S1: *Temporal comparisons: experimental or natural perturbations* for references), and annual rates of change (i.e., per year) were estimated for 63 study-treatment combinations; rates of change for all treatments (including controls) were quantified in our analyses. Natural and experimental treatments included changes due to warming, eutrophication, fire, grazing, restoration, severe storms or other disturbances, and kelp removal. Taxonomic groups included algae, plants, invertebrates, fish, birds, and mammals.

### Spatial comparisons: natural environmental variation

We combined two existing compilations of data to examine spatial patterns of biodiversity change in relatively natural environmental contexts. The CESTES database (Jeliazkov et al. 2019) contains assemblage data from studies that sampled species at multiple sites (it also includes information on traits and environment that we do not use here); we removed studies with explicit human impacts identified as an environmental feature, and our analysis included 19 studies that sampled terrestrial, freshwater and marine assemblages from a number of taxonomic groups (birds, plants, insects, macroinvertebrates, fishes and mammals). McGill (2011b) compiled datasets with two or more local assemblages containing species abundance data; we removed studies documenting disturbances and other perturbations, resulting in 32 studies being retained. From the combined 51 studies, those with many sites were randomly subsampled down to ten sites so that they did not dominate the results. Within each study, an arbitrary site was assigned as the ‘reference’ site, and change was quantified between every site and the reference within studies; our analysis included a total of 356 spatial comparisons.

### Spatial comparisons: anthropogenic perturbations

To quantify spatial differences in biodiversity due to anthropogenic land use, we used the PREDICTS database (Hudson et al. 2017). We used the 2016 release of the database (downloaded from https://data.nhm.ac.uk/dataset/the-2016-release-of-the-predicts-database on 10_th_ July 2020). We limited our analyses to studies with abundance data for individuals, and those with known land use categories (primary vegetation, mature secondary vegetation, intermediate secondary vegetation, plantation forest, cropland, pasture, and urban); studies where land use was not recorded were omitted. This resulted in 237 combinations of source ID and study (some sources had multiple studies, denoted SS in the database), and 418 estimates of change relative to the reference land use (primary vegetation) category.

### Statistical models

To estimate changes in the different metrics whilst accounting for expected correlations between them, we fit multivariate multilevel models to the data. Similar to the way univariate multilevel (also called hierarchical or mixed-effects) models fit to a single response can allow varying (also called random) intercepts and slopes to be correlated, this approach estimates changes in all components simultaneously whilst allowing for (and estimating) correlations between them.

Response distributions for all metrics were chosen to ensure changes were estimated on similar measurement scales, and because all metrics take only positive values, log response scales were used for all components.

For the *Temporal comparisons: natural environmental variation* data, total abundance (*N*) was fit with a model that assumed a lognormal distribution and identity link function, and Poisson distributions with log link functions were fit to *S*_n_, *S*_PIE_ and *S*; Poisson distributions were chosen for *S*_n_ and *S*_PIE_ values rounded to integers based on visual assessments that showed lognormal models fit to raw *S*_n_ and *S*_PIE_ values greatly underpredicted the density of ones in the data. For the *Temporal comparisons: experimental or natural perturbations* data, *S* was no longer an integer value after standardizing sampling effort, and all metrics were fit with models that assumed lognormal distributions and identity link functions. Both spatial data sets were fit with models that assumed lognormal distributions and identity link functions for total abundance (*N*), rarefied richness (*S*_n_) and evenness (*S*_PIE_), and a Poisson distribution and log-link function for species richness (*S*).

The *Temporal comparisons: natural environmental variation* models included non-varying intercepts and slopes for year, and varying intercepts and slopes for studies and cells for all responses. To allow for correlations between changes in the different responses, varying study- and cell-level parameters for all responses were drawn from a single multivariate normal distribution for each level (i.e., one for studies, one for cells; see Appendix S1: *Temporal comparisons: natural environmental variation* for equations). The models fit to the *Temporal comparisons: experimental or natural perturbations* data similarly included non-varying intercepts and slopes for year, and had varying intercepts for study, site and block fitted separately for each response. For these data, correlations between changes in the different responses were modeled by drawing varying intercepts and slopes for each combination of treatment and study for all responses from a single multivariate normal distribution (see Appendix S1: *Temporal comparisons: experimental or natural perturbations* for equations).

The models fit to the *Spatial comparisons: natural environmental variation* data included non-varying intercepts for data source (i.e., CESTES and McGill). Correlations between the different responses were modeled by assuming varying intercepts and slopes (representing the reference site and departures for all other sites from the reference, respectively) for each study and response came from a single multivariate normal distribution; over-dispersion in the richness response was modeled using an observation-level varying intercept (see Appendix S1: *Spatial comparisons: natural environmental variation* for equations). Models fit to the *Spatial comparisons: anthropogenic perturbations* data included non-varying intercepts and slopes (representing the reference [primary vegetation] category and departures from the reference for each land use category, respectively), and varying intercepts for sites and blocks were modeled separately for each response. Correlations between changes in the different responses were modeled by assuming that varying intercepts and slopes (as per the non-varying intercepts and slopes) for each combination of source and study and each response came from a single multivariate normal distribution (see Appendix S1: *Spatial comparisons: anthropogenic perturbations* for equations).

All statistical models were estimated using the Hamiltonian Monte Carlo (HMC) sampler Stan (Carpenter et al. 2017), and coded using the ‘brms’ package (Burkner 2017). Details of all model specifications, and the iterations and warmup periods are provided in the Appendix, as are the priors (which were weakly regularizing). Visual inspection of the HMC chains and model diagnostics (Rhat < 1.05) showed good mixing of chains and convergence, and model adequacy assessed visually using posterior predictive checks showed that the models were able to make predictions similar to the empirical data (see Appendix Figure: S1-4). Code for all analyses is available at https://github.com/sablowes/MulticomponentBioChange, and will be archived following publication.

## Results

Temporal changes in perturbed environments had the highest percentage of assemblages with at least one component trend (Δ*N,* Δ*S*_n_, Δ*S*_PIE_, or Δ*S*) that differed from zero (44%), followed by spatial comparisons across land use categories (29%), temporal changes (21%) and spatial comparisons in naturally varying environments (12%). Component changes that differed from zero showed broadly similar patterns across datasets, with one exception: trends differing from zero for multiple components were less common for spatial comparisons between assemblages in naturally varying environments (Appendix S1: Figure S5).

Pairwise relationships between changing components were typically positive (i.e., had the same sign), though exceptions to this general pattern were found for all data sources (Figure 3). For assemblages where Δ*N* and Δ*S*_PIE_ had the same sign (though not necessarily differing statistically from zero), richness changes were typically larger in magnitude (Figure 3). In contrast, assemblages where Δ*N* and Δ*S*_PIE_ had opposing signs typically exhibited changes in richness that were smaller in magnitude (Figure 3). This tendency for countervailing changes in abundance and evenness to constrain richness changes was most apparent for spatial comparisons between different land use categories (Figure 3j-l), and there was a high proportion of assemblages that were growing in size (Δ*N* > 0) but with declining species richness (Δ*S* < 0; Figure 3j), associated with declining evenness (Δ*S*_PIE_ < 0).

**Figure 3:**
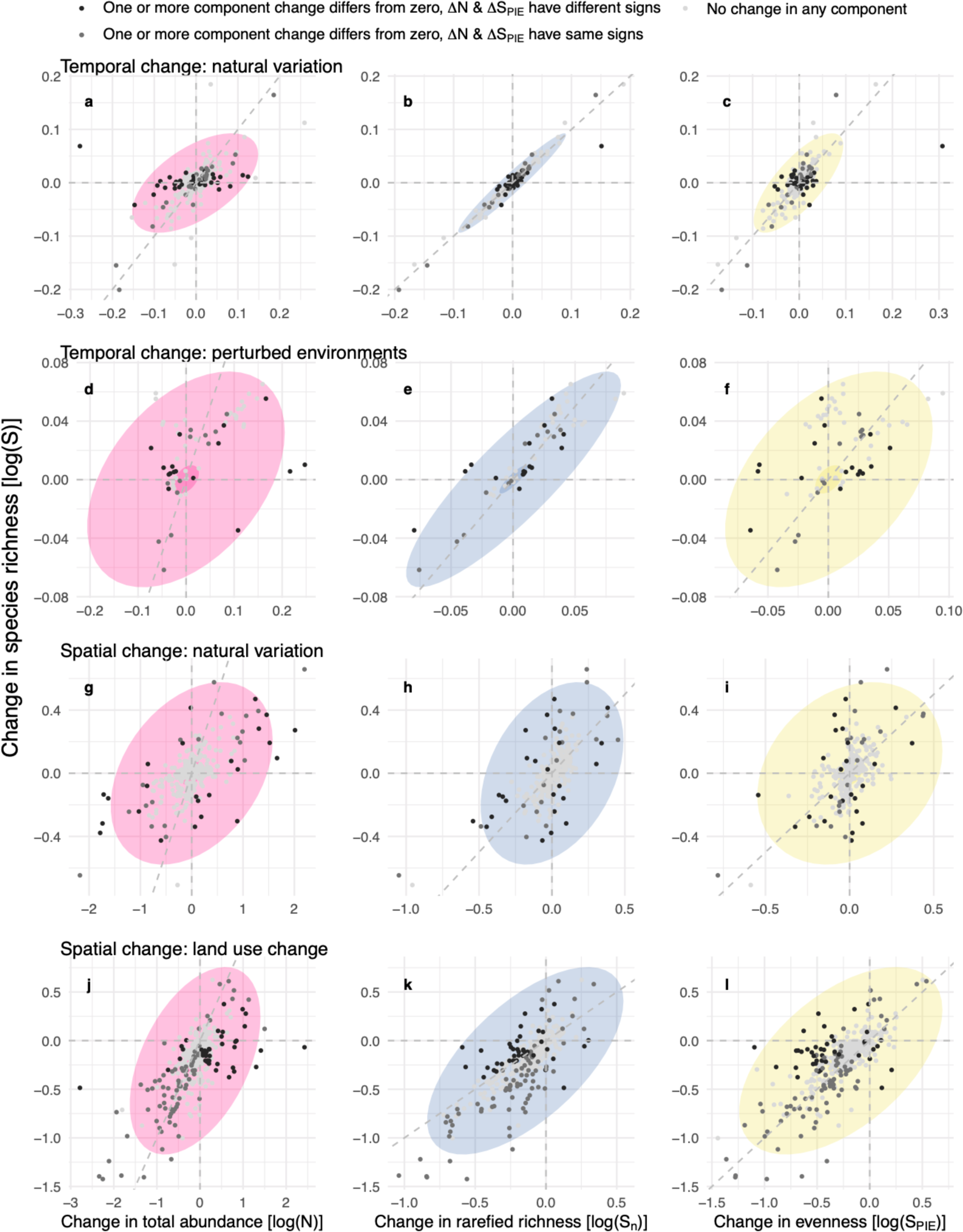
Empirical relationships between four components of local diversity change. Change in species richness as a function of changes in the numbers of individuals (left column), rarefied richness (middle column), and evenness (right column) for (a-c) study-level estimates of temporal changes in naturally varying environments; (d-f) estimates of temporal change for combinations of study and treatment in perturbed environments; (g-i) estimates of spatial changes within studies from an arbitrary reference site along natural environmental gradients; and, (j-l) estimates of spatial change within studies between primary vegetation and different land use categories. Colored concentration ellipses show the confidence interval (5 and 95%) of the posterior distributions. Dotted grey lines are x and y = 0, and x = y. See Appendix S1: Figure S6 for remaining pairwise relationships. Scale of x- and y-axes vary between panels; one estimate with Δlog(*N*) = -1.79, Δlog(*S*) = -3.77, Δlog(*S*_n_) = -3.23, Δlog(*S*_PIE_) = -3.21, removed from (j-l) for clarity.

The strongest relationships were found for components changing through time, and relationships between richness and changes in the SAD -- rarefied richness (Figs. 3b, e) and evenness (Figs. 3c, f; Figs. 4a, b) -- were stronger than those between changes in richness and total abundance (Figs. 3a, c; Figs. 4a, b). Spatial comparisons had generally weak relationships overall. No strong relationships between changing components emerged for comparisons in natural environments (Figs. 3g-i; Fig. 4c), and only weakly positive relationships between changes in abundance, evenness and richness were found in comparisons between primary vegetation and different land use categories (Figure 3j-l, Figure 4d).

**Figure 4:**
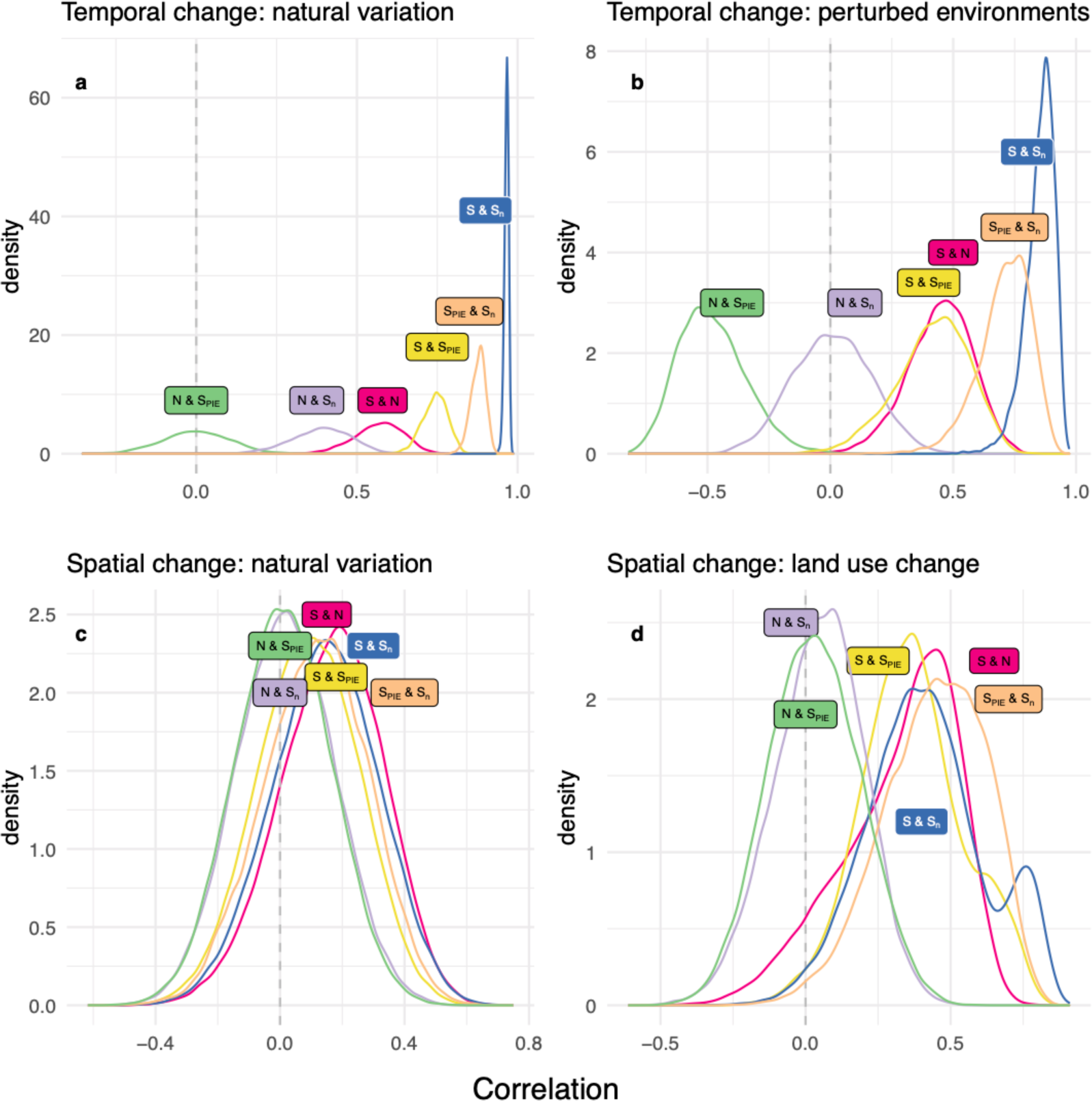
Component correlations among studies within each data source. Density plots for the posterior distribution of pairwise correlations between component changes for (a) temporal comparison in naturally varying environments, (b) temporal comparisons in perturbed environments, (c) spatial comparisons along natural gradients, and (d) spatial comparisons between different land use categories. Correlations estimated separately for sites and land use categories relative to the references were combined on (c) and (d).

Temporal changes in naturally varying assemblages were roughly centered on zero for all metrics (Figure 3a-c). Across assemblages, altered numbers of individuals and species richness changes had a moderately positive relationship (Figure 4a), weakened predominantly by assemblages that had opposing abundance and evenness relationships (Figure 3a). In contrast, relationships between changes in species richness and rarefied richness, and between richness changes and evenness changes were strong (Figure 4a). Assemblages in perturbed environments had slightly positive temporal trends on average in all components (Figure 3d-f). Across assemblages, Δ*S* and Δ*N* (Figure 3d, 4b) and Δ*S* and Δ*S*_PIE_ (Figure 3f, 4b) had relatively weak positive relationships, whereas Δ*S* and Δ*S*_n_ (Figure 3e, 4b) showed a strong positive relationship.

Spatial comparisons in naturally varying environments exhibited highly heterogeneous patterns of change centered around zero for all metrics (Figure 3g-i). Decoupled component changes meant that relationships between them were generally absent or weak across assemblages (Figure 4c). Spatial comparisons between assemblages in primary vegetation versus those in different land use categories were also highly heterogeneous, though there were typically fewer individuals, less even assemblages and fewer species relative to primary vegetation (Figure 3j-l). Across assemblages, land use change was typically associated with relatively weak positive relationships between changes in the components of local diversity (Figure 4d).

## Discussion

Our conceptual overview using individual-based rarefaction curves clearly shows how the expectation of positive pairwise relationships between changes in abundance, evenness and richness arises. If curves stretch or contract, we expect positive relationships between changes in total abundance and richness. Similarly, if curves flex upwards or downwards, positive relationships between changes in evenness and richness are expected. Rarefaction curves also show how contrasting signs of changes in abundance and evenness can strongly determine the magnitude of richness changes, and determine whether positive relationships between changes in richness and the other components (abundance and evenness) are likely. Both these predictions were generally well supported by our empirical analyses. Relationships between changes in abundance, evenness and richness were generally positive, and richness changes were typically greater for assemblages with strictly positive pairwise relationships. Countervailing changes in total abundance and evenness, where found, often constrained the magnitude of changes in species richness, and acted to weaken relationships between Δ*N* and Δ*S*, and Δ*S*_PIE_ and Δ*S*. Spatial comparisons had the most heterogeneous relationships between changes in abundance, evenness, and richness, and in relatively natural environments changes were sufficiently decoupled that no strong relationships emerged across assemblages. In contrast, strong positive correlations between temporal changes in species richness (Δ*S*) and changes in metrics associated with altered SADs (Δ*S*_n_, Δ*S*_PIE_) emerged across assemblages. These temporal results show strong support for the prediction that variation in relative abundances can dominate local variation in biodiversity (Chapin et al. 2000), even when human impacts are less direct.

### Variation in assemblage size does not dominate local diversity change

Overall, only ∼2% of assemblages in this study (22 of 1125) had changes consistent with a strong ‘sampling’ effect on changes in species richness (i.e., Δ*N* & Δ*S* having the same sign, and being the only changes different from zero). This finding complements existing evidence showing that despite many tests, empirical evidence for the more-individuals hypothesis (Srivastava & Lawton 1998) remains equivocal (Storch et al 2018, Vagle & McCain 2020).

While both (species-level) population variability and variation associated with sampling (Vagle & McCain 2020) likely contribute to the weak response of species richness to variation in the total number of individuals, our results are broadly consistent with previous syntheses showing that broad-scale spatial variation in richness was rarely driven simply by variation in the numbers of individuals (Currie et al. 2004, Storch et al 2018). Our results indicate that variation in local diversity, through time or from site-to-site, is not due to changes in assemblage size passively sampling more or less from a static SAD. Instead, we show that variation in local biodiversity can be strongly influenced by changes to species’ relative abundances. These changes can be occurring at multiple scales (Hillebrand et al. 2008, Blowes et al. 2020), and could reflect altered local environmental conditions (e.g., altered resource or habitat availability and diversity, eutrophication, local harvest or exploitation), or changes at broader scales that alter the species pool (via species additions or subtractions).

Our general result showing that variation in the total abundance of an assemblage through time or space is often decoupled from changes in metrics of biodiversity such as species richness also cautions against making “apples to oranges” comparisons in the context of quantifying biodiversity change. For example, some estimates of change are based on either population-level abundance (e.g., Living Planet Index, WWF 2020), or assemblage-level abundance (e.g., insect declines; Wagner 2000, van Klink et al. 2020), whereas other change estimates are based on patterns of the number or identity of species present (e.g., Dornelas et al. 2014, Newbold et al. 2015). Our results show that assuming abundance and richness changes are strongly correlated will often be an oversimplification. Moreover, the importance of altered relative abundances for local biodiversity variation means that biodiversity change estimates will frequently depend on whether changes in species relative abundances influence the metrics used (see e.g., Antão et al. 2020b).

### Contrasting component relationships between temporal changes versus spatial comparisons of biodiversity

Relationships between changing components of biodiversity showed strikingly different patterns between temporal changes and spatial comparisons. Moreover, these differences were generally greater than those found between naturally-varying and perturbed assemblages, for either temporal changes or spatial comparisons.

Pairwise relationships between changes in abundance, evenness and richness were typically weak for spatial comparisons. Decoupling was greatest, and pairwise relationships weakest, for changes between sites experiencing relatively natural environmental variation. However, given our simple conceptual framework shows that some degree of interdependence cannot be avoided, we caution against overinterpreting the relative independence of these component changes, and further analyses examining component change relationships along continuous spatial gradients are warranted. Indeed, evenness and richness are never numerically independent (Jost 2010), and the weak overall relationship between changes in richness and evenness for these data was in part due to assemblages with countervailing changes in abundance and evenness. Most importantly, these highly variable component changes further emphasize the need for a holistic approach to quantifying biodiversity change (Chase et al. 2018, Avolio et al. 2021).

Our prediction that the signs of changes in abundance and evenness can strongly determine the magnitude of richness changes was most evident in the spatial contrasts between primary vegetation and other land use categories (Newbold et al. 2015, 2020). Assemblages with the greatest declines in abundance and evenness had the greatest richness declines. In contrast, when abundance and evenness changes had opposing signs, richness changes were tempered. Indeed, countervailing abundance and evenness changes were frequently associated with components other than species richness (i.e., Δ*N*, Δ*S*_n_, and/or Δ*S*_PIE_) having a trend that differed from zero across all data sources (Appendix S1: Table S1). This highlights that even apparently decoupled or weakly correlated component changes have interdependencies that can remain important determinants of observed changes.

In contrast to assemblage change between sites, there was strong coupling between species richness and SAD changes through time. In particular, the strength of the relationship between Δ*S*_n_ and Δ*S* resulted in estimates of change being similar for most assemblage time series in relatively natural environments (Figure 3b). In some cases, this occurred despite countervailing changes in total abundance and evenness (Figure 3a, b). For assemblages where abundance and evenness changed in the same direction, similar estimates of Δ*S*_n_ and Δ*S* indicate that abundance changes were occurring along a relatively flat region of the individual-based rarefaction curve. This shows that changes to the total number of individuals need not strongly influence species richness, even where signs are the same and they have a positive relationship. The strong association between richness changes and altered relative abundances has important implications for examining causes and/or consequences of biodiversity change (Hillebrand et al. 2008, Crowder et al. 2010). Even where the expected positive relationships between abundance, evenness and richness are found, we can more fully understand assemblage changes when all component changes are examined simultaneously.

While both approaches, time series and spatial comparisons (or space-for-time substitutions), have contributed to our understanding of biodiversity change, the relative merits of each for our understanding of ecological dynamics has not been discussed much (Adler et al. 2020). The largely decoupled component changes found here for spatial comparisons suggest that too much focus on average changes across assemblages, such as those in total abundance or in species richness, risks masking highly heterogeneous changes occurring within assemblages in multiple components. Moreover, decoupled, heterogeneous component changes complicate using spatial comparisons to infer temporal changes. The smaller effect sizes found here for time series indicates greater inertia compared to site-to-site variation. More generally, the strong role of changes to the SAD for variation in local biodiversity suggests that deepening our understanding of altered patterns of relative abundance across scales represents an important direction for future theoretical and empirical work. Here our focus has been on numerical relationships between component changes, and using process-based models (e.g., Thompson et al. 2020) to examine how altered metacommunity dynamics impact patterns of relative abundance across scales could help our understanding how different processes impact component relationships. Similarly, empirical studies could ask whether local environmental changes are affecting evenness, or if changes occurring at broader spatial scales are impacting the size of species pool and the regional SAD?

## Conclusions

We found strong correlations between changes in the SAD and species richness changes through time, whereas relationships between abundance and richness changes for both temporal and spatial diversity variation were generally weak. Our findings confirm that altered species relative abundances, and/or changes to the size of the species pool, often strongly influence local diversity change (Chapin et al. 2010), even where human impacts are less direct. However, our results also reinforce cautions against examining changes to any one component of biodiversity change in isolation (e.g., Wilsey et al. 2005, Chase et al. 2018, Avolio et al. 2021).

To be most useful, quantifying (co)variation in the different components of biodiversity needs to be done coherently. Individual-based rarefaction curves and associated metrics can provide an intuitive and illustrative characterization of relationships among changing components of local biodiversity. Whilst ecologists are increasingly looking beyond species richness to quantify biodiversity change (e.g., Dornelas et al. 2014, Hillebrand et al. 2018), different components of biodiversity and its change within assemblages are most often analyzed independently, and frequently with metrics lacking conceptual unity. Conceptually and empirically, our results emphasize that changes to the most frequently quantified aspects of biodiversity, including changes to the numbers of individuals, and the relative abundance and total number of species are highly interdependent. Examining how within-assemblage component changes covary with potential drivers could reveal insights masked by independent aggregate estimates of change across assemblages, and provide new information for understanding biodiversity change in the Anthropocene.

## Acknowledgements

SAB thanks the Biodiversity Synthesis group at iDiv for contructive feedback at various stages of the project. SAB, TE, AS, and JMC were supported by the German Centre for Integrative Biodiversity Research (iDiv) Halle-Jena-Leipzig, funded by the German Research Foundation (FZT 118).

## Appendix S1

**Table S1:**
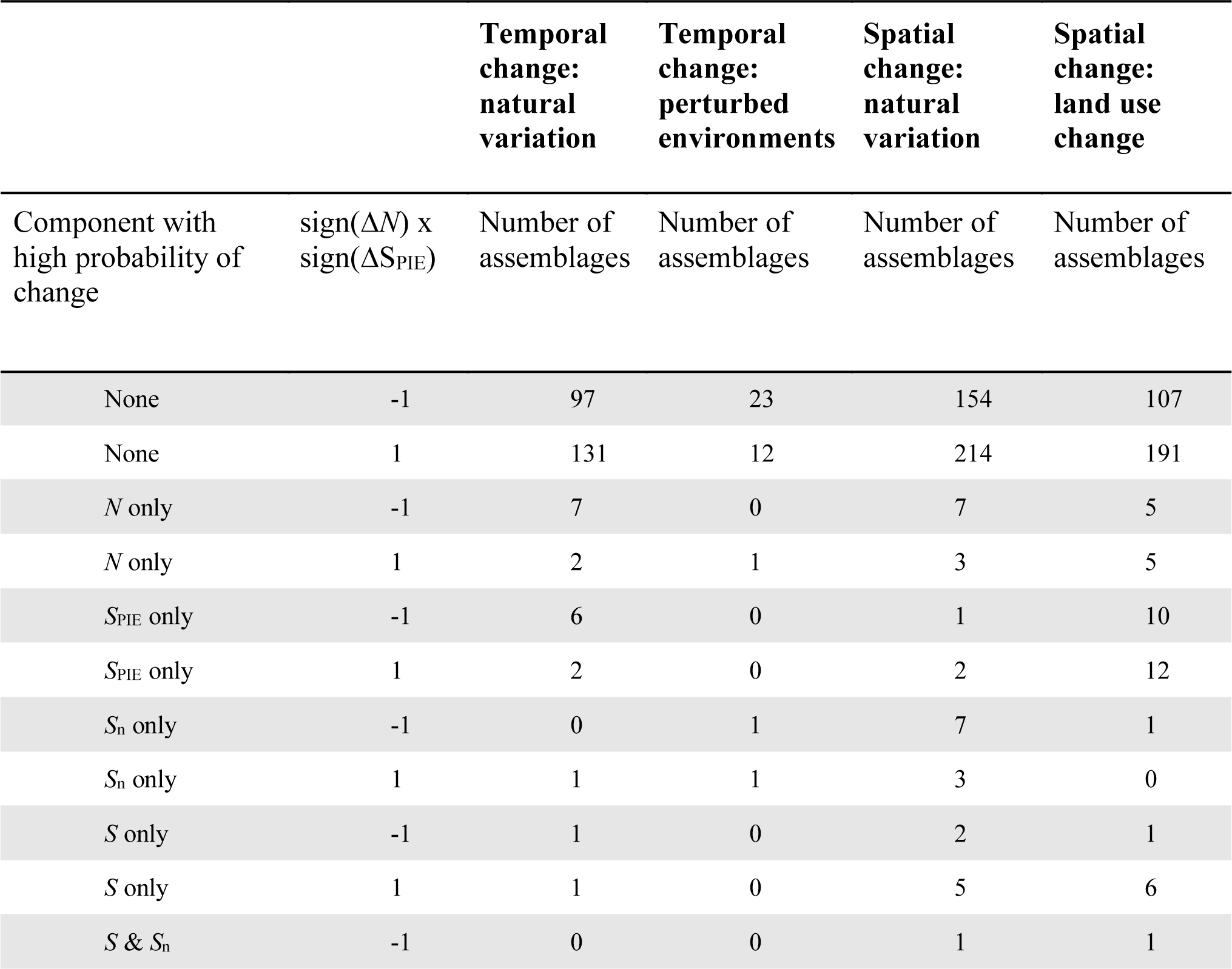
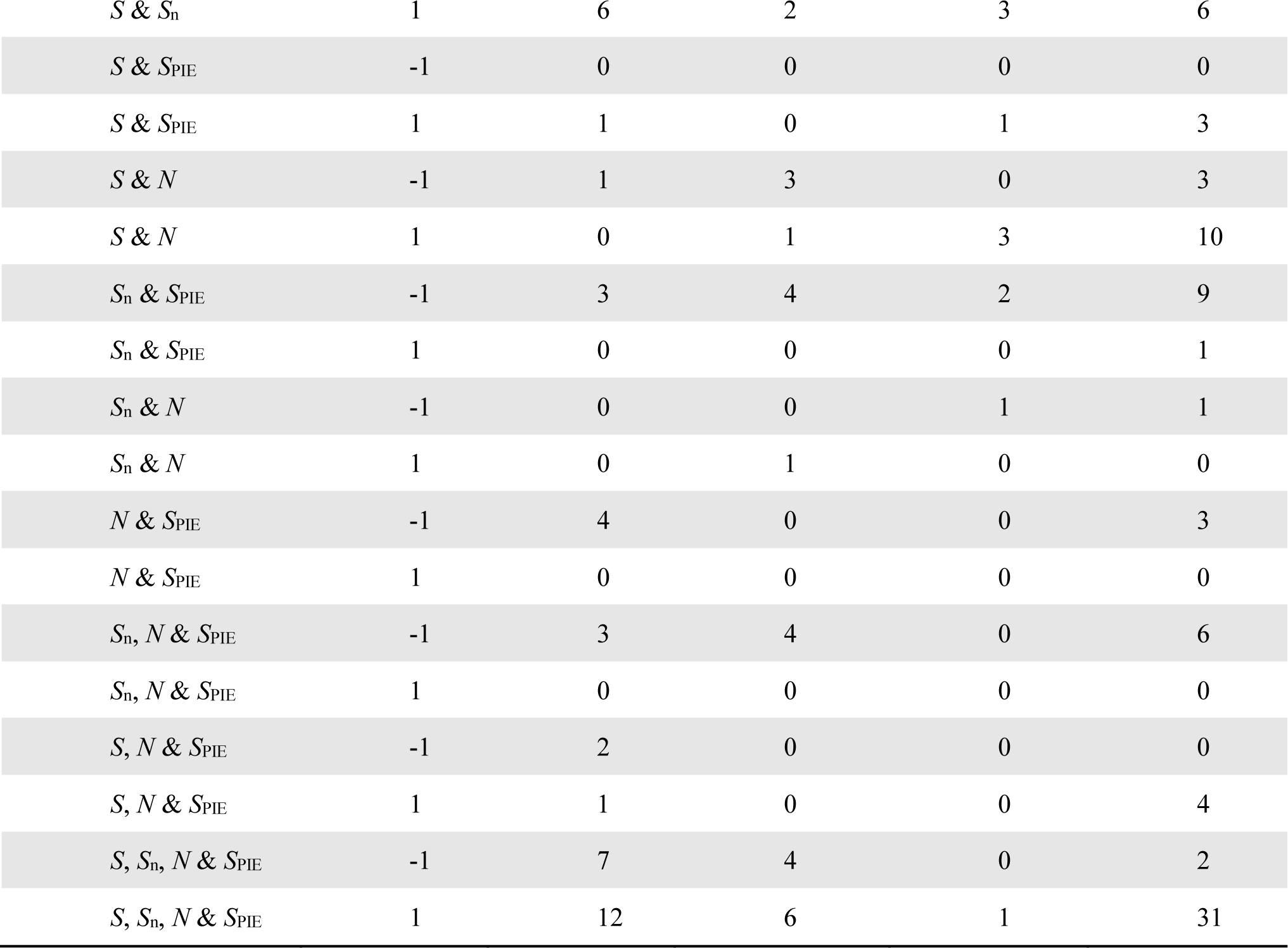
Countervailing changes in abundance (Δ*N*) and evenness (Δ*S*_PIE_) often result in components other than richness changing. Component(s) with high probability of change as per Figure 2 (i.e., 90% CI does not overlap zero); each row shows the number of assemblages with either countervailing [sign(Δ*N*) x sign(ΔS_PIE_) = -1] or abundance and evenness changes with the same direction [sign(Δ*N*) x sign(ΔS_PIE_) = 1] for the different data sources.

### Temporal comparisons: natural environmental variation

We fit a model that assumed a lognormal distribution for total abundance (*N*), and poisson distributions (with log link functions) for species richness (S), rarefied richness (*S*_n_) and evenness (*S*_PIE_). We ran the model with four chains for 2000 iterations, with 1000 used as warmup. The model took the following form:

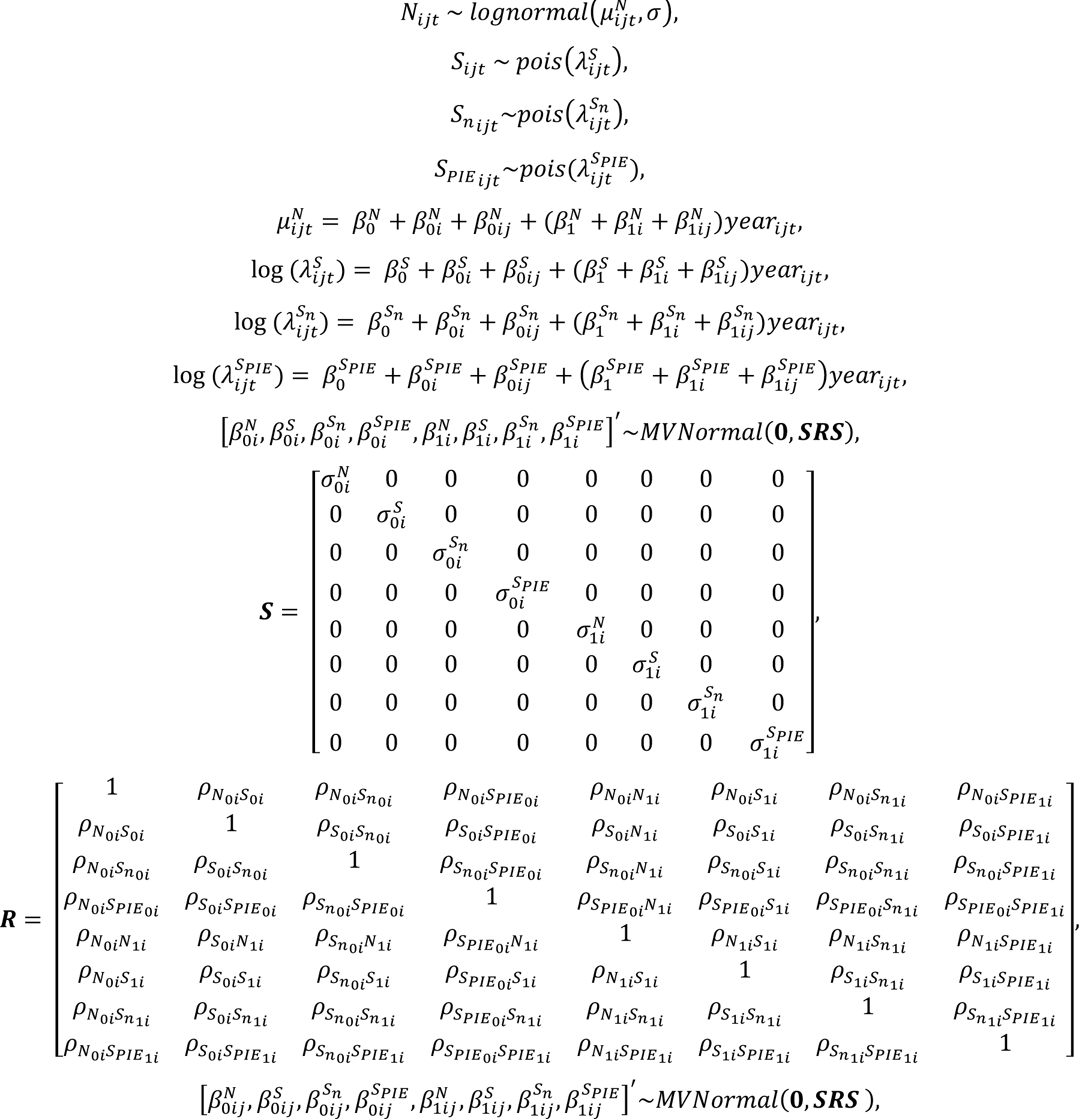

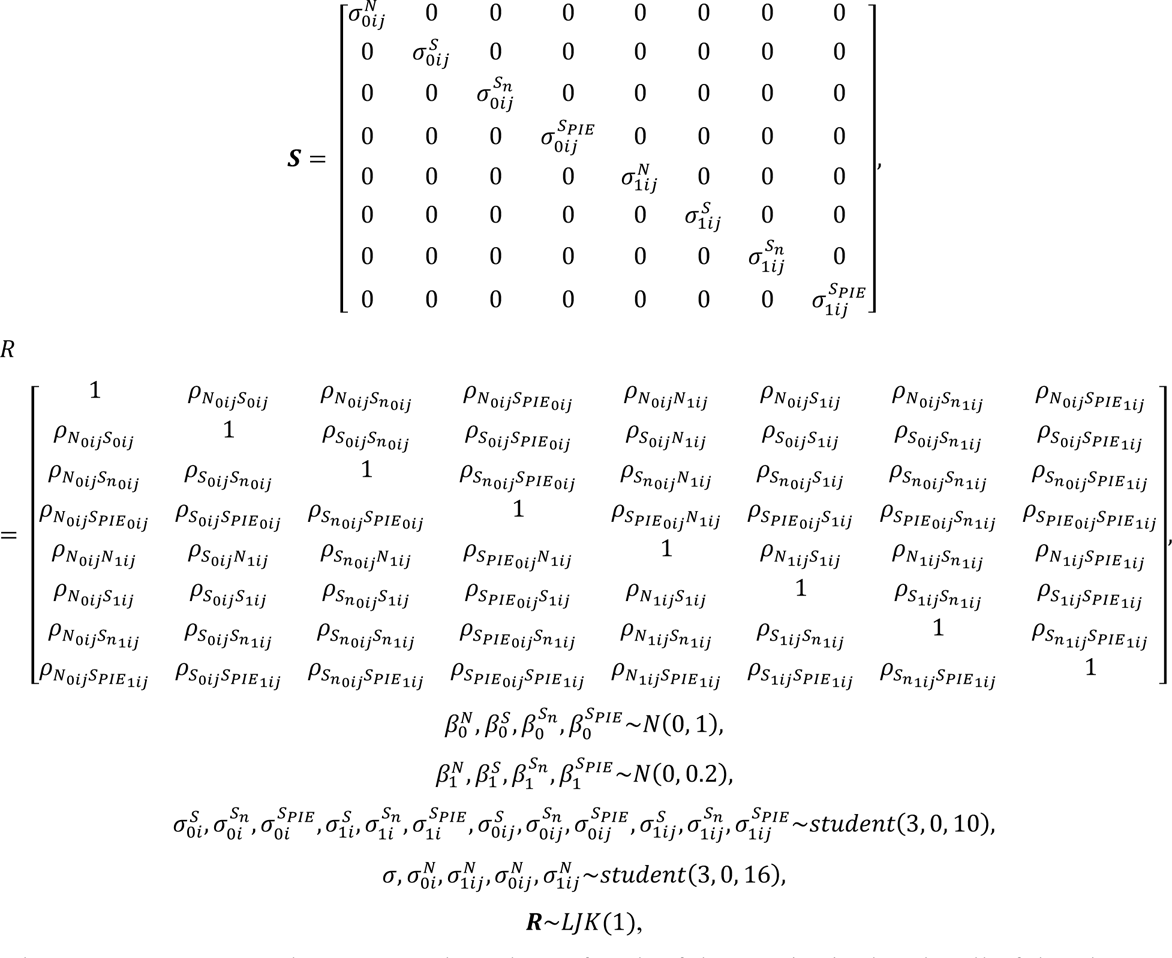

where *N*_ijt_, *S*_ijtt_ *S*_nijkt_, and *S*_*PIE*_ijkt__ are the values of each of the metrics in the *j*th cell of the *i*th study in year *t*, and year_ijt_ is the time in years. β_0_ and β_1_ (with the respective superscripts for each metric) represent the non-varying intercepts and slopes, respectively. β_0i_ and β_1i_ (with the respective superscripts for each metric) represent the varying intercepts and slopes for study-level departures from β_0_ and β_1_, respectively. β_0ij_ and β_1ij_ (with the respective superscripts for each metric) represent the varying intercepts and slopes for cell-level departures from β_0_ and β_1_, respectively. The covariance matrices of each multivariate normal distribution for varying effects (one each for the study- and cell-level departures) are parameterised in terms of a correlation matrix **R** and two matrices **S** with diagonal elements 2 (superscripts for metrics, subscripts denote intercept (0), slopes (1) and level: studies *i* and cells *j*).

**Figure S1:**
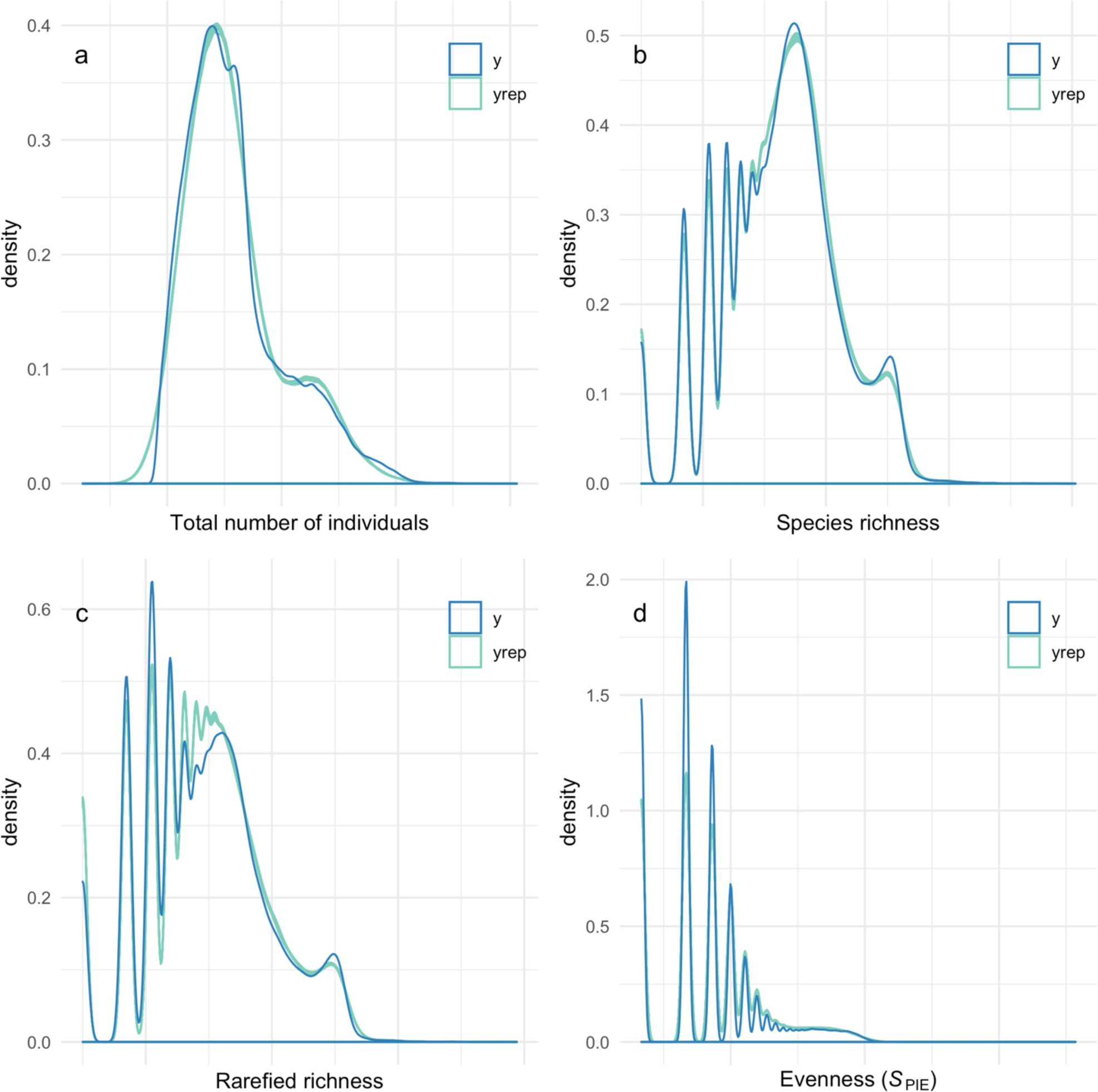
Posterior predictive checks for the model fit to the temporal data in naturally varying environments. (a) Total numbers of individuals, (b) species richness, (c) rarefied richness (Sn) and (d) evenness (SPIE). Each panel shows the density of the data (y) and ten draws of the posterior distribution (yrep).

### Temporal comparisons: experimental or natural perturbations

We fit a model that assumed lognormal distributions for all response variables (*N, S*, *S*_n_, *S*_PIE_). Our focus is on the treatment-level variation in the metrics, and we created a covariate that was the concatenation of study and treatment to this end. Additionally, as some studies had blocks within sites, and others did not, we created a new variable that was the concatenation of site and block, where those without block had no unique identifier. We ran the model with four chains for 3000 iterations, with 1500 used as warmup. The model took the following form:

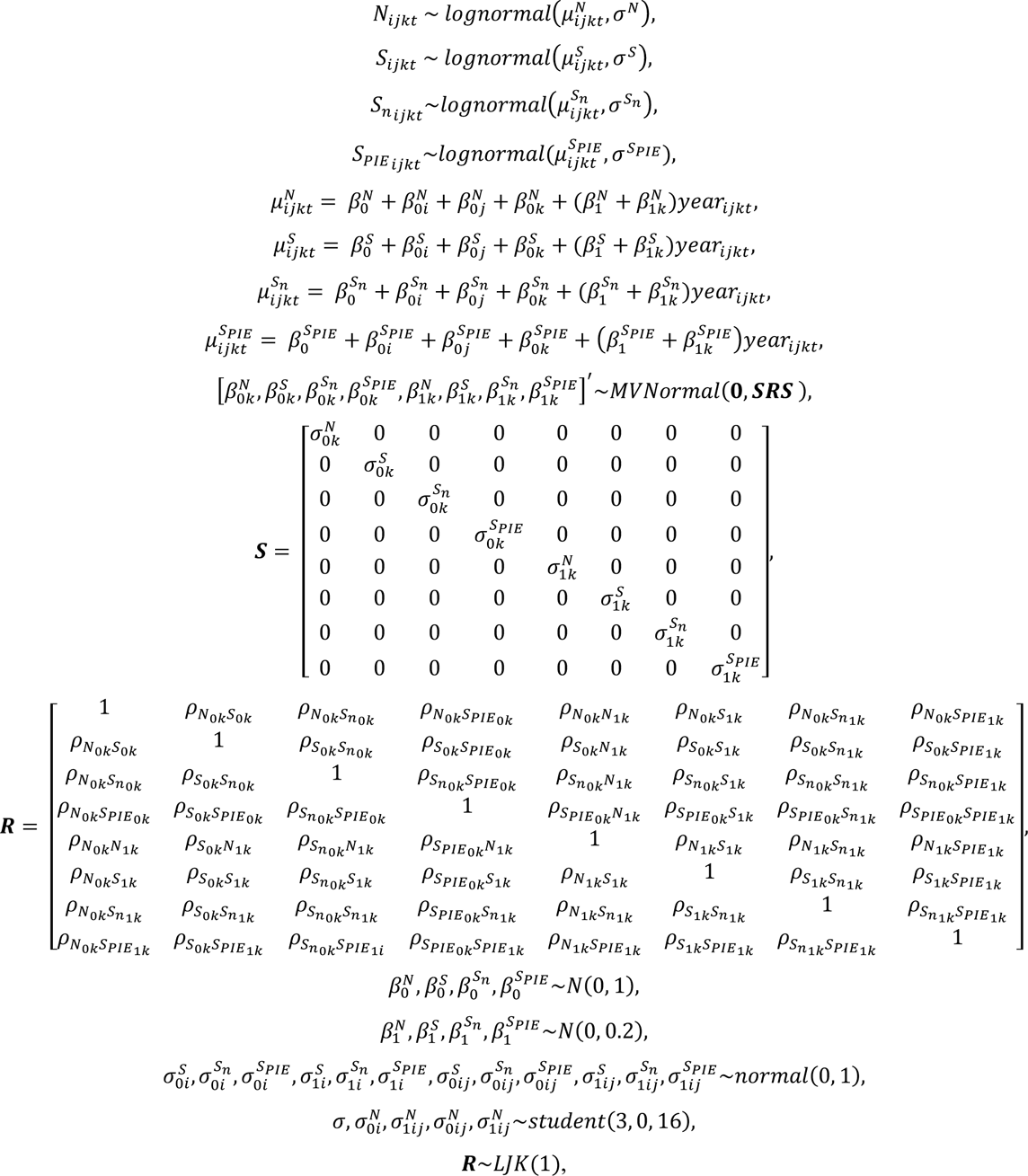

where *N*_ijkt_, *S*_ijkt_, *S*_*nij*_, and *S*_*PIE*_ij__ are the values of each of the metrics in the kth study-treatment combination, for the *j*th study-block combination of the *i*th study in year *t*, and year_ijkt_ is the time in years. β_0_ and β_1_ (with the respective superscripts for each metric) represent the non-varying intercepts and slopes, respectively. β_0i_ (with the respective superscripts for each metric) represent the varying intercepts study-level departures from β_0_, and β_0j_ (with the respective superscripts for each metric) represent the varying intercepts study-block level departures from β_0_. β_0k_ and β_1k_ (with the respective superscripts for each metric) represent the varying intercepts and slopes for study-treatment departures from β_0_ and β_1_, respectively. The covariance matrix of the multivariate normal distribution for varying study-treatment effects were parameterised in terms of a correlation matrix **R** and two matrices **S** with diagonal elements 2 (superscripts for metrics, subscripts denote intercept (0), slopes (1) and study-treatment combination, *k*).

**Figure S2:**
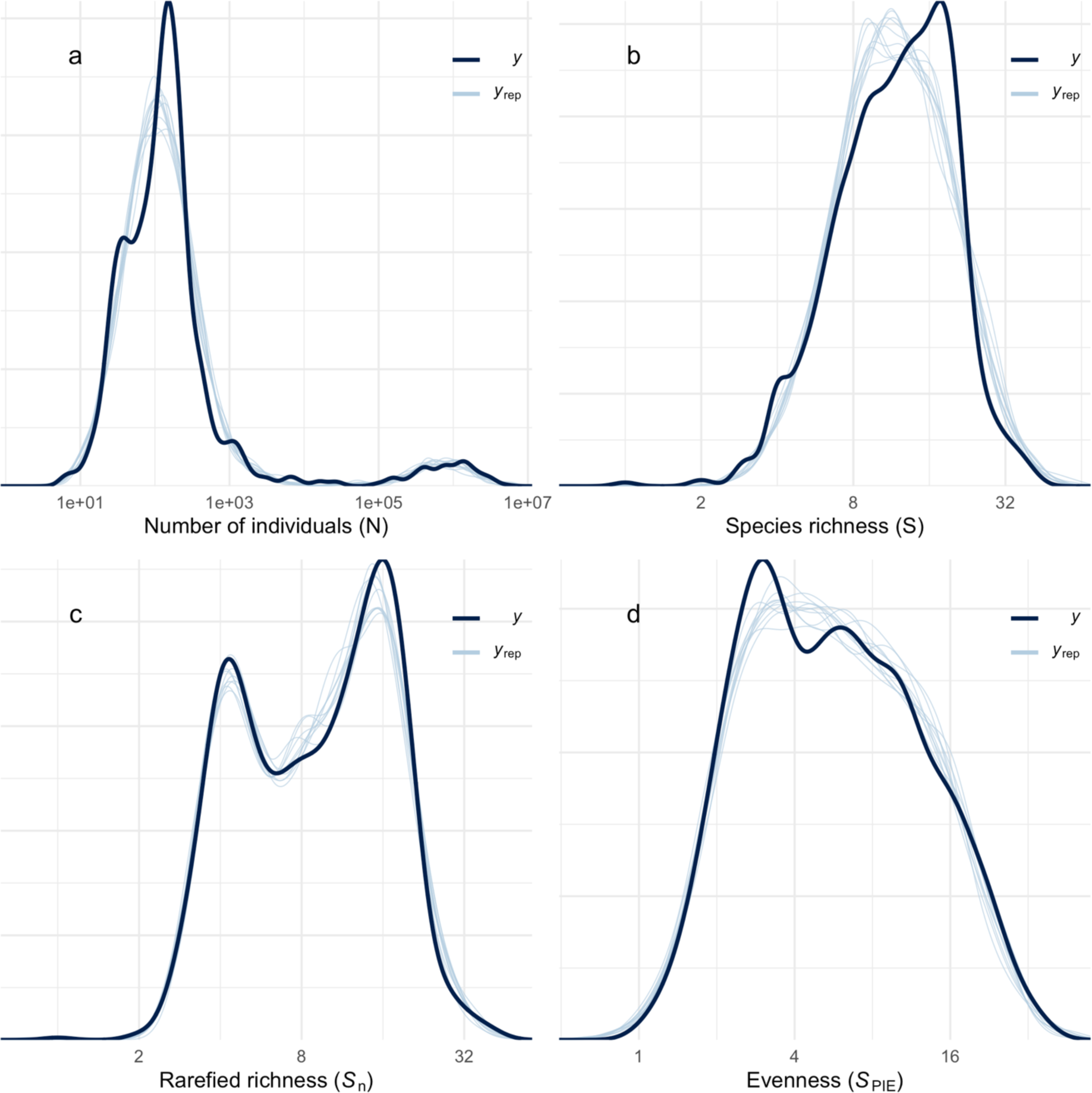
Posterior predictive checks for the model fit to the temporal data in perturbed environments. (a) Total numbers of individuals, (b) species richness, (c) rarefied richness (Sn) and (d) evenness (SPIE). ). Each panel shows the density of the data (y) and ten draws of the posterior distribution (yrep).

### Spatial comparisons: natural environmental variation

We fit a model that assumed lognormal distributions for *N*, *S*_n_, and *S*_PIE_, and a Poisson distribution (and log link function) for *S.* We fit the model with four chains and 4000 iterations, with 2000 as warmup. The model took the following form:

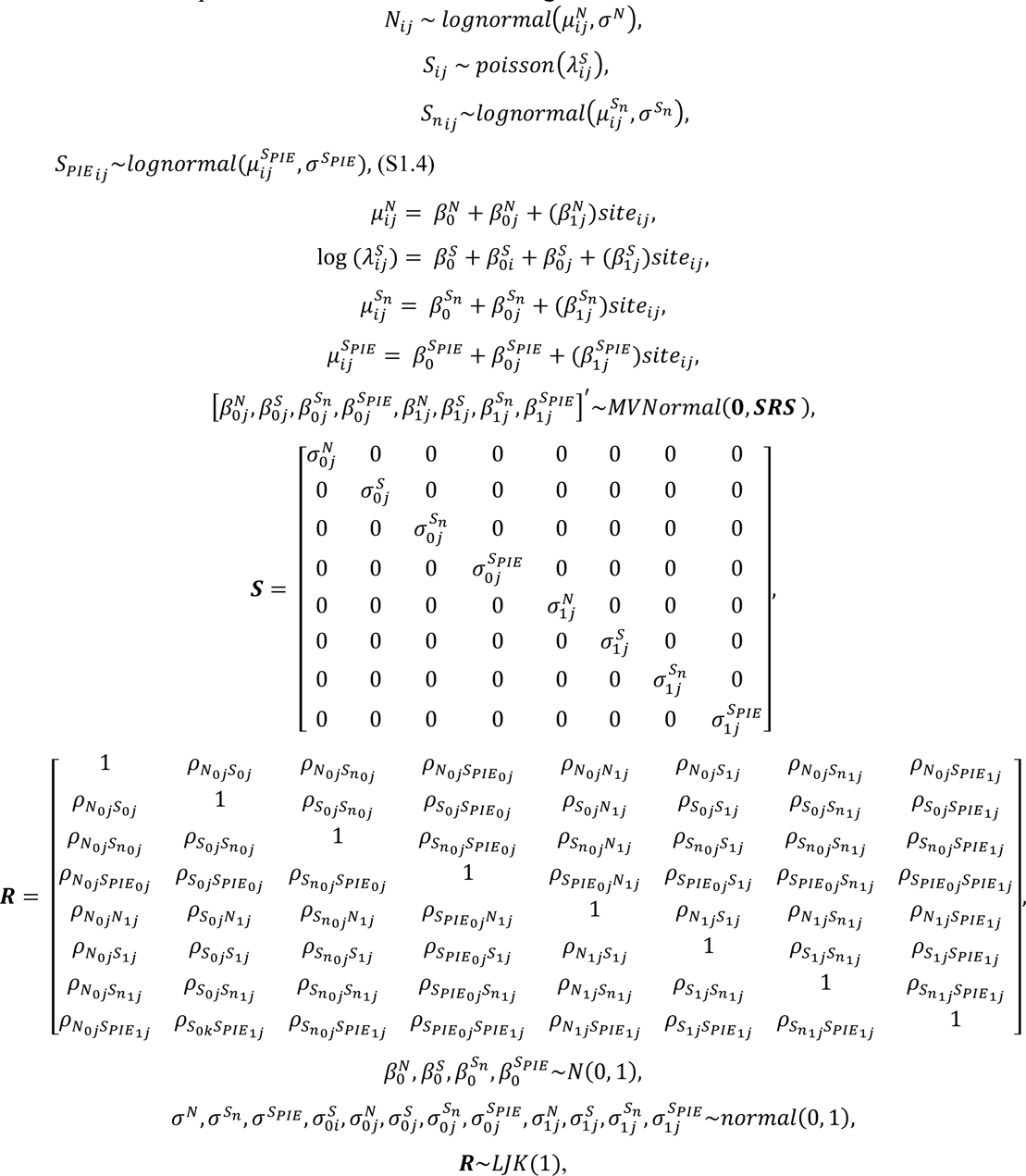

where *N*_ijt_, *S*_ijtt_ *S*_nijkt_, and *S*_*PIE*_ijkt__, are the values of each of the metrics for the *i*th observation in the

*j*th study-site combination, and sitβ_ij_ is site identifier. β_0_ (with the respective superscripts for each metric) represent the non-varying intercepts for each data source (one each for CESTES and

McGill), respectively. β_0i_^*S*^ is an observation level varying intercept to model overdispersion in *S*.

E0j and E1j (with the respective superscripts for each metric) represent the varying intercepts (reference sites for each study) and slopes (departures for all other sites). The covariance matrix of the multivariate normal distribution for varying study-site effects were parameterised in terms of a correlation matrix **R** and two matrices **S** with diagonal elements 2 (superscripts for metrics, subscripts denote intercept (0), slopes (1)).

**Figure S3:**
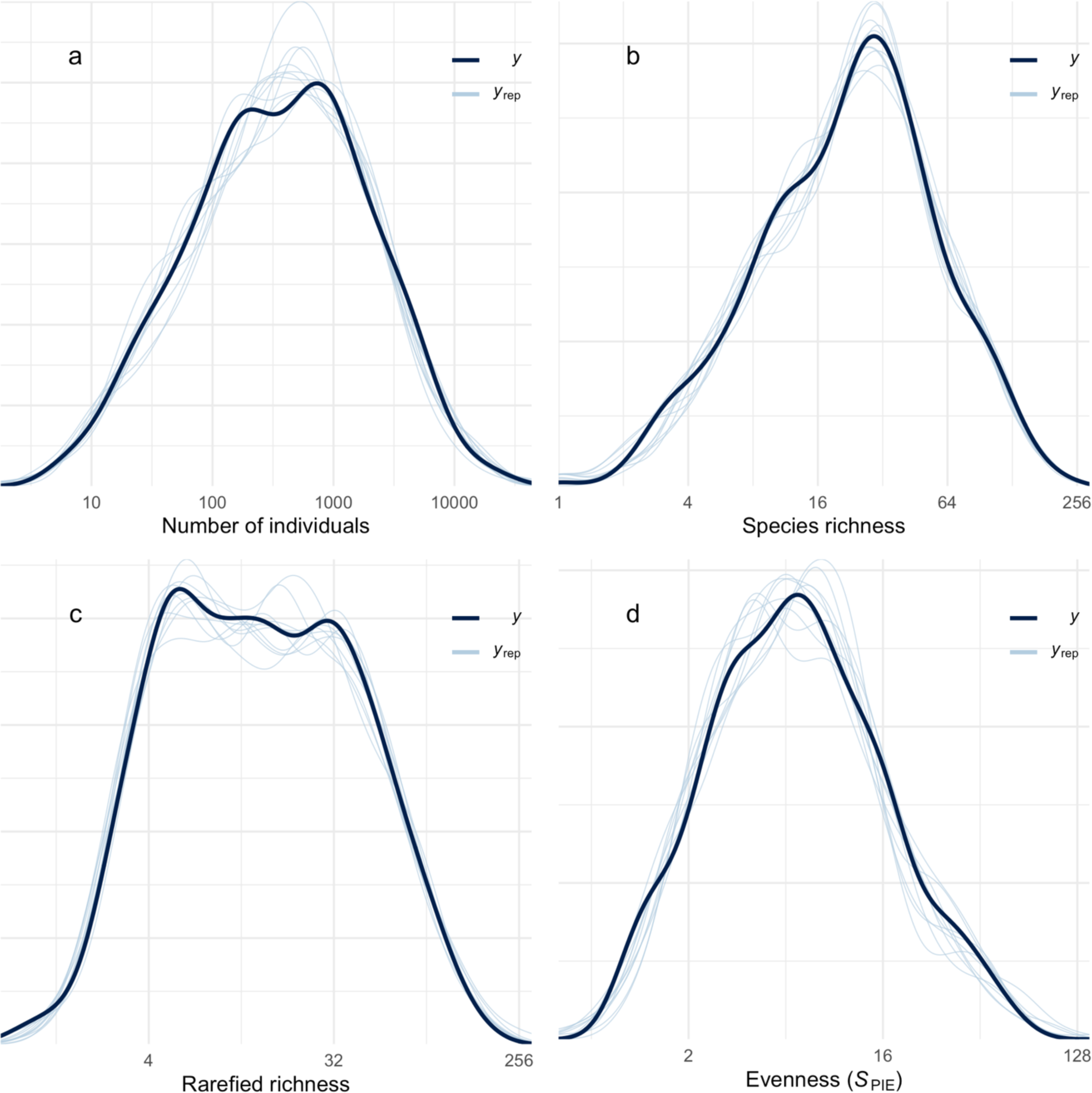
Posterior predictive checks for the model fit to the spatial data in natural environments. (a) Total numbers of individuals, (b) species richness, (c) rarefied richness (Sn) and (d) evenness (SPIE). Each panel shows the density of the data (y) and ten draws of the posterior distribution (yrep).

### Spatial comparisons: anthropogenic perturbations

We fit a model that assumed lognormal distributions for *N*, *S*_n_, and *S*_PIE_, and a Poisson distribution (and log link function) for *S*. We fit the model with four chains and 4000 iterations, with 2000 used as warmup. The model took the following form:

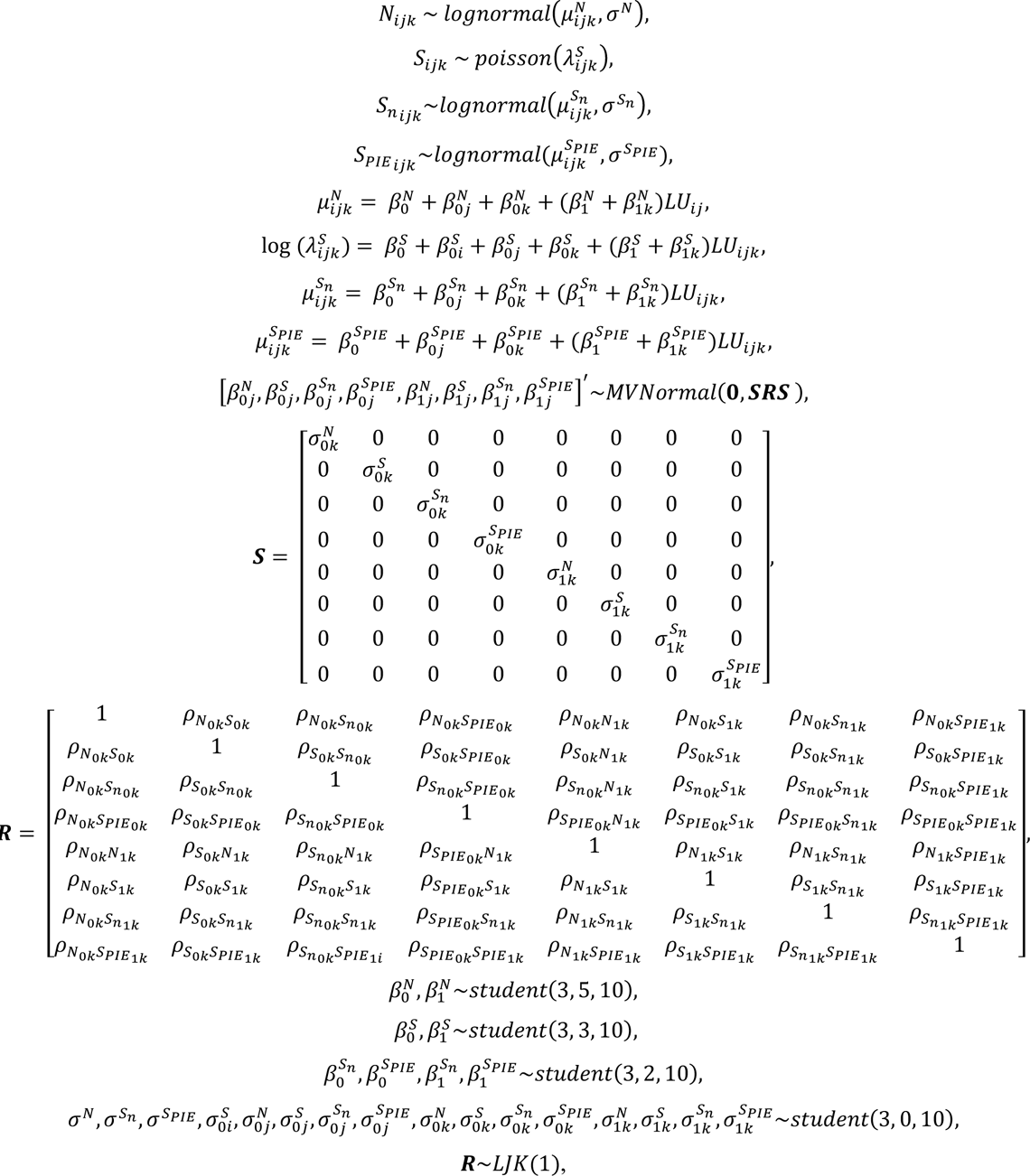

where *N*_ijt_, *S*_ijtt_ *S*_nijkt_, and *S*_*PIE*_ijkt__ are the values of each of the metrics for the *i*th concatenation of source ID, study number, block and site number (denoted SSBS in the database), in the *j*th concatenation of source ID, study number, block (denoted SSB in the database), of the *k*th combination of source ID and study (denoted SS in the database), and *LU* is an identifier for land use types (other than primary vegetation, that was fit as the intercept). β_0_ (with the respective superscripts for each metric) represent the non-varying intercepts for reference land use category (i.e., primary vegetation). β_0i_^*s*^ is a site-level varying intercept to model overdispersion in *S*. β_0j_ (with the respective superscripts for each metric) is a varying intercept for blocks. β_0k_ and β_1k_ (with the respective superscripts for each metric) represent the varying intercepts (reference sites for each study) and slopes (departures for all other sites). The covariance matrix of the multivariate normal distribution for varying study-site effects were parameterised in terms of a correlation matrix **R** and two matrices **S** with diagonal elements 2 (superscripts for metrics, subscripts denote intercept (0), slopes (1)).

**Figure S4:**
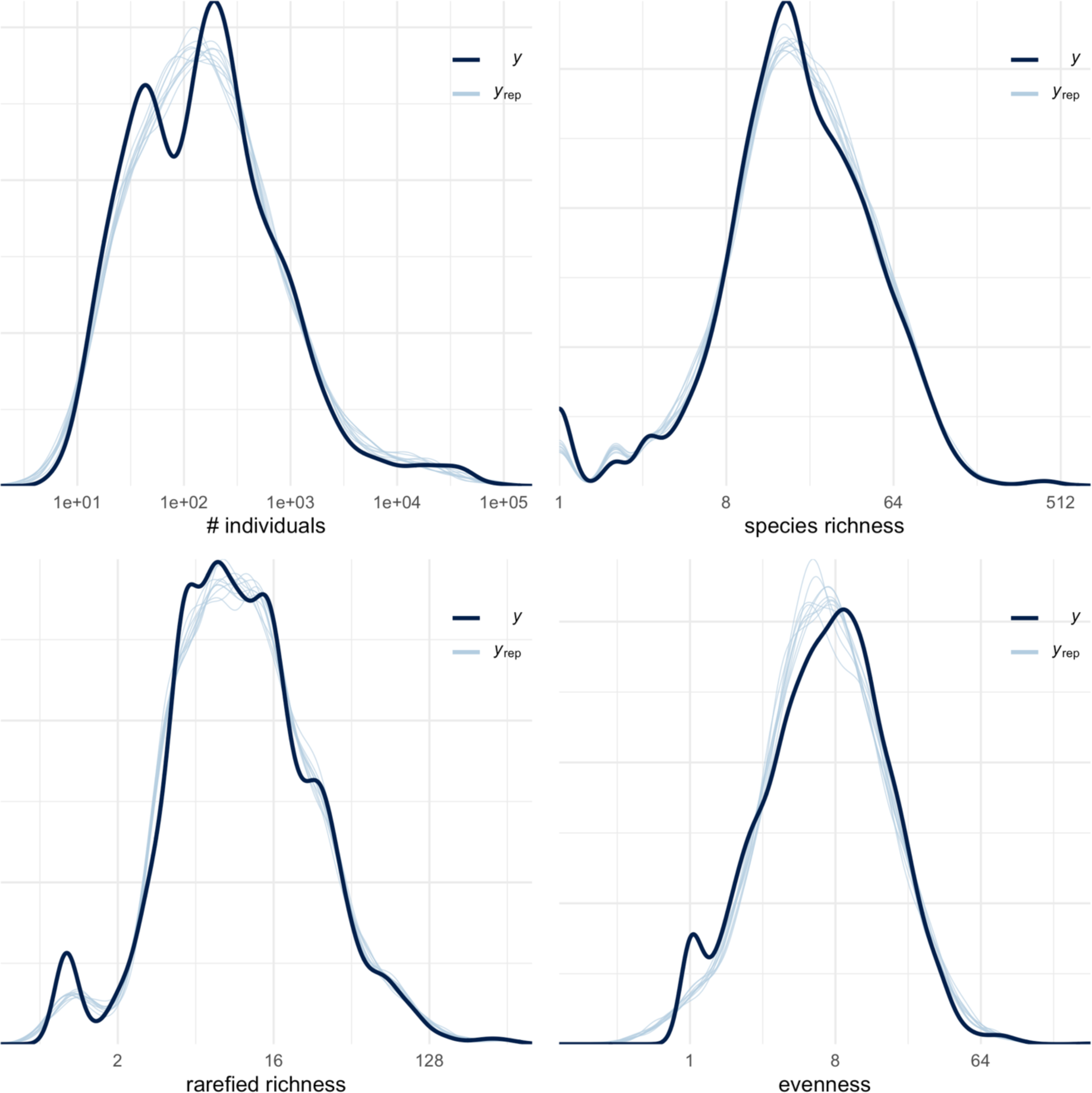
Posterior predictive checks for the model fit to the spatial data in anthropogenically perturbed environments. (a) Total numbers of individuals, (b) species richness, (c) rarefied richness (Sn) and (d) evenness (SPIE). Each panel shows the density of the data (y) and ten draws of the posterior distribution (yrep).

**Figure S5:**
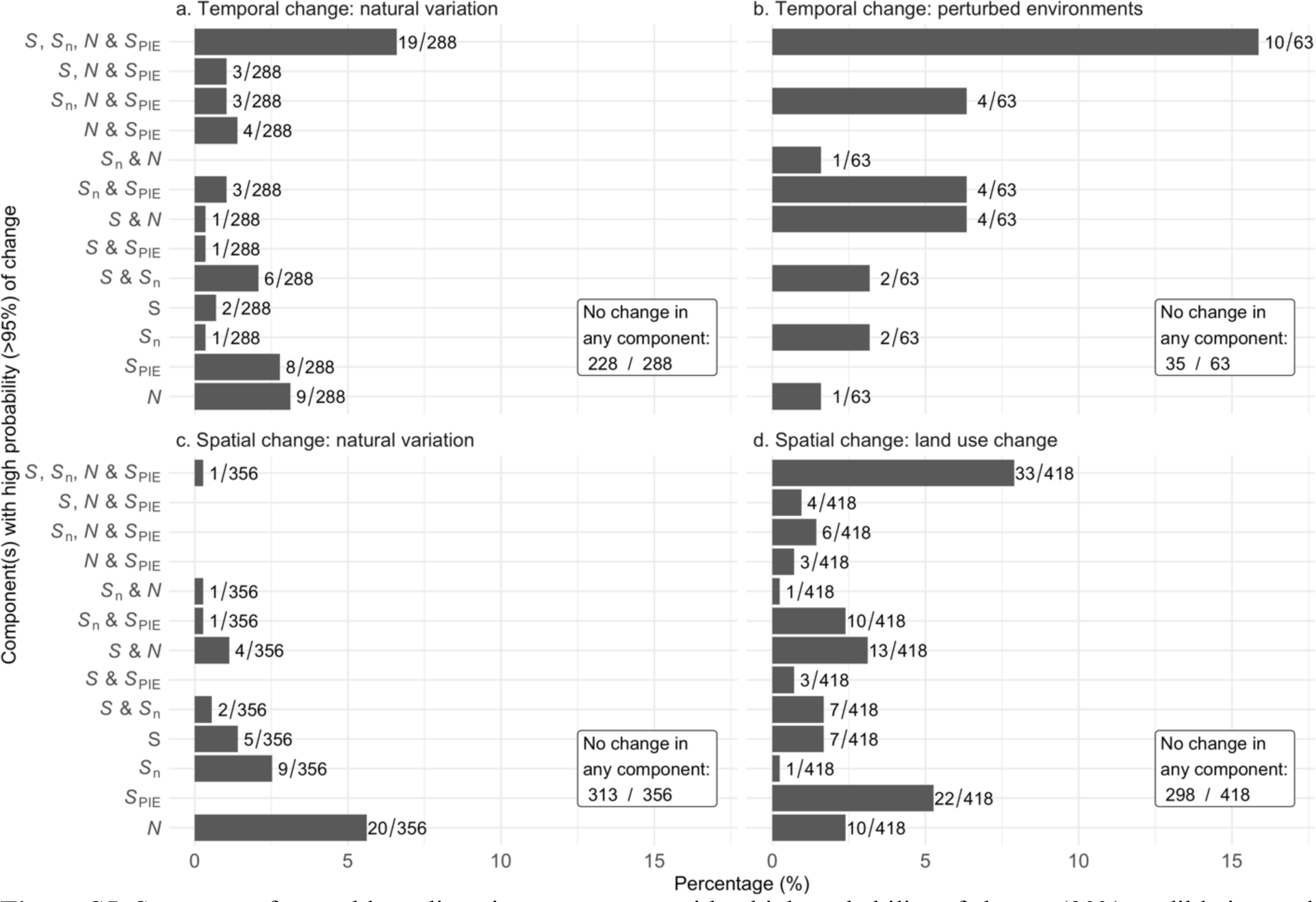
Summary of assemblage diversity components with a high probability of change (90% credible interval did not overlap zero).(a) temporal changes in naturally varying environments, (b) temporal changes in perturbed environments, (c) spatial changes relative to an arbitrary reference, (d) spatial changes relative to primary vegetation. Assemblages with no component changes different from zero are reported as insets for clarity. Metric abbreviations: total number of individuals (*N*), expected number of species for n individuals (*S*n), numbers equivalent transformation of the Probability of Interspecific Encounter (*S*PIE), and total species richness (*S*). Number following each bar is the count of assemblages for that category.

**Figure S6:**
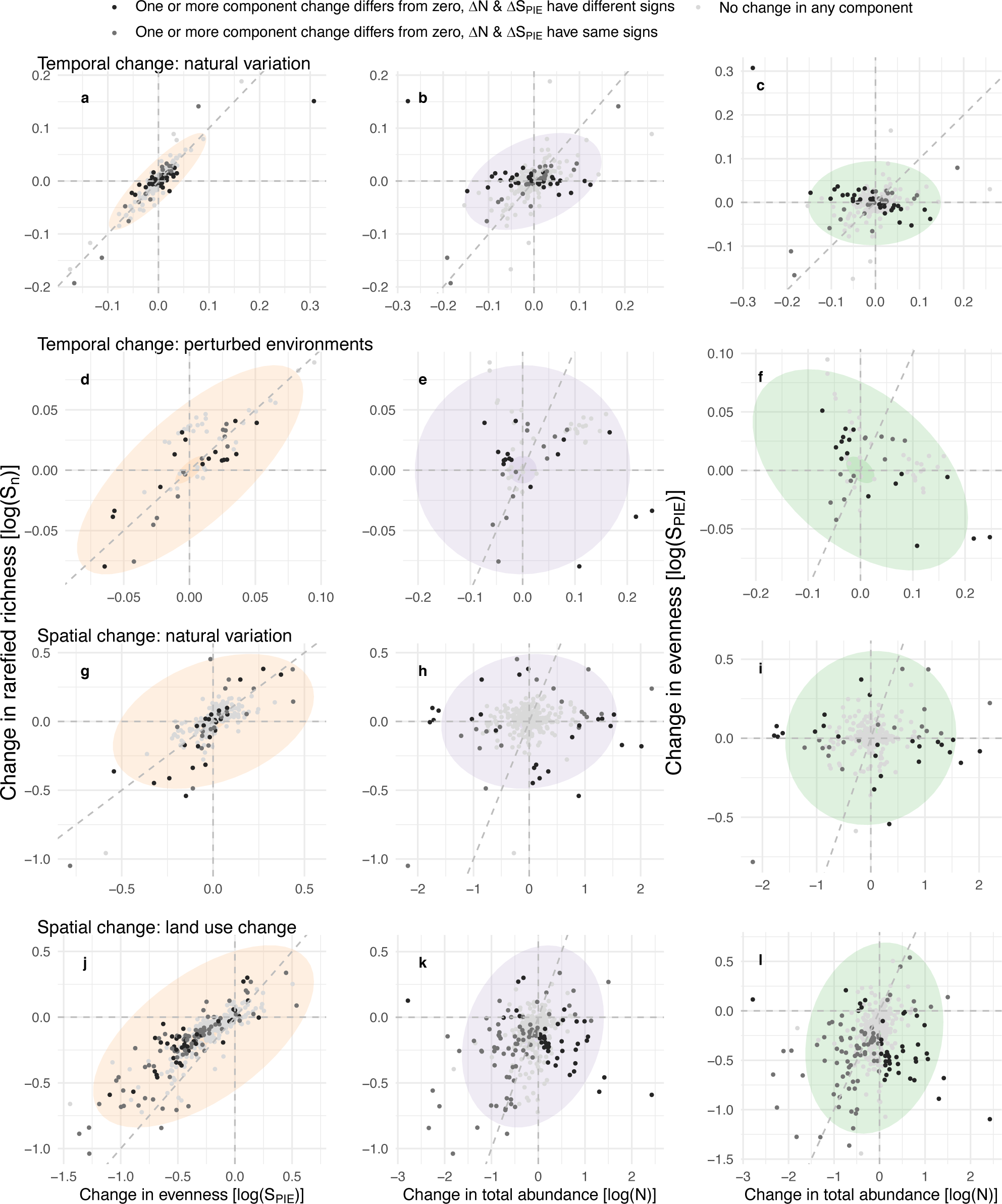
Relationships between four components of local diversity change (not shown on Figure 3). Change in rarefied richness as a function of changes in evenness (left), change in rarefied richness as a function of changes in total abundance (middle) and evenness (right) for (a-c) study-level estimates of temporal changes in naturally varying environments; (d-f) estimates of temporal change for combinations of study and treatment in perturbed environments; (c) estimates of spatial changes within studies from an arbitrary reference site along natural environmental gradients; and, (d) estimates of spatial change within studies between primary vegetation and different land use categories. Coloured concentration ellipses show 10% increments (5 – 95%) of the posterior distributions. Dotted grey lines are x = y = 0, and x = y for visual reference. NB: Scale of x- and y-axes vary between data sources; one estimate with ΔN = -1.79, ΔS = -3.77, ΔSn = -3.23, ΔSPIE = -3.21, removed from (j-l) for clarity.

